# Optimal Control Costs of Brain State Transitions in Linear Stochastic Systems

**DOI:** 10.1101/2022.05.01.490252

**Authors:** Shunsuke Kamiya, Genji Kawakita, Shuntaro Sasai, Jun Kitazono, Masafumi Oizumi

**Affiliations:** Graduate School of Arts and Sciences, The University of Tokyo; Araya, Inc.

## Abstract

The brain is a system that performs numerous functions by controlling its states. Quantifying the cost of this control is essential as it reveals how the brain can be controlled based on the minimization of the control cost, and which brain regions are most important to the optimal control of transitions. Despite its great potential, the current control paradigm in neuroscience uses a deterministic framework and is therefore unable to consider stochasticity, severely limiting its application to neural data. Here, to resolve this limitation, we propose a novel framework for the evaluation of control costs based on a linear stochastic model. Following our previous work, we quantified the optimal control cost as the minimal Kullback-Leibler divergence between the uncontrolled and controlled processes. In the linear model, we established an analytical expression for minimal cost and showed that we can decompose it into the cost for controlling the mean and covariance of brain activity. To evaluate the utility of our novel framework, we examined the significant brain regions in the optimal control of transitions from the resting state to seven cognitive task states in human whole-brain imaging data. We found that, in realizing the different transitions, the lower visual areas commonly played a significant role in controlling the means, while the posterior cingulate cortex commonly played a significant role in controlling the covariances.

**Significance Statement:** The brain performs many cognitive functions by controlling its states. Quantifying the cost of this control is essential as it reveals how the brain can be optimally controlled in terms of the cost, and which brain regions are most important to the optimal control of transitions. Here, we built a novel framework to quantify control cost that takes account of stochasticity of neural activity, which is ignored in previous studies. We established the analytical expression of the stochastic control cost, which enables us to compute the cost in high-dimensional neural data. We identified the significant brain regions for the optimal control in cognitive tasks in human whole-brain imaging data.

## 1 Introduction

The brain is a highly complex dynamical network that flexibly transitions to various states to execute a myriad of functions [1, 2, 3]. In this regard, the brain can be considered a system that modulates its internal states to desired states, in accordance with the function the individual needs to perform [4, 5]. Among the many transitions that bring the system into the various states it requires, some state transitions are more difficult to control than others, depending on the dynamical properties of the neuronal systems. In other words, controlling transitions to some states incurs greater “costs” than controlling transitions to others. Providing a theoretical framework for quantifying transition costs, or control costs, is important for evaluating the difficulty of the shifts between various brain states, and possibly in explaining cognitive loads [6, 7], sleep-awake differences [8], habituation of cognitive tasks [9], and psychiatric disorders [7] with a quantifiable measure. Therefore, the development of such a framework for quantifying control cost in the brain is a vital topic in neuroscience.

A rigorous and promising framework to assess control cost was provided by an approach using control theory, which was first introduced in neuroscience in a pioneering work by Gu et al. [5]. Control theory provides theoretical tools for investigating the dynamical properties of complex systems [10, 11], and its application to neuroscience is opening new doors to mechanistically explaining neural behaviors from brain structures [7, 9, 12, 13, 14, 15]. However, despite being a strong approach, this framework does not take account of an important property of neural activity: it neglects noise or stochasticity in neural systems. Since neural noises are known to be ubiquitous in the brain and to play critical roles in information processing [16, 17], overlooking the stochasticity of neural systems may result in an inaccurate estimation of the control costs. In this study, we propose a novel framework to quantify control costs in linear stochastic neural systems. This framework takes advantage of both the linear control theoretic framework [5, 9, 13] and the control cost proposed in our previous work [6]. That is, we modeled brain dynamics using linear stochastic differential equations, and defined the control cost as the KL divergence between the uncontrolled and controlled processes (Fig. 1) as we did in [6]. Thanks to the linearity, we can obtain an analytical expression of the control cost in the stochastic system. Furthermore, as we include a control input term in our model, we can identify those brain regions playing a significant role in the control of state transitions.

**Figure 1:**
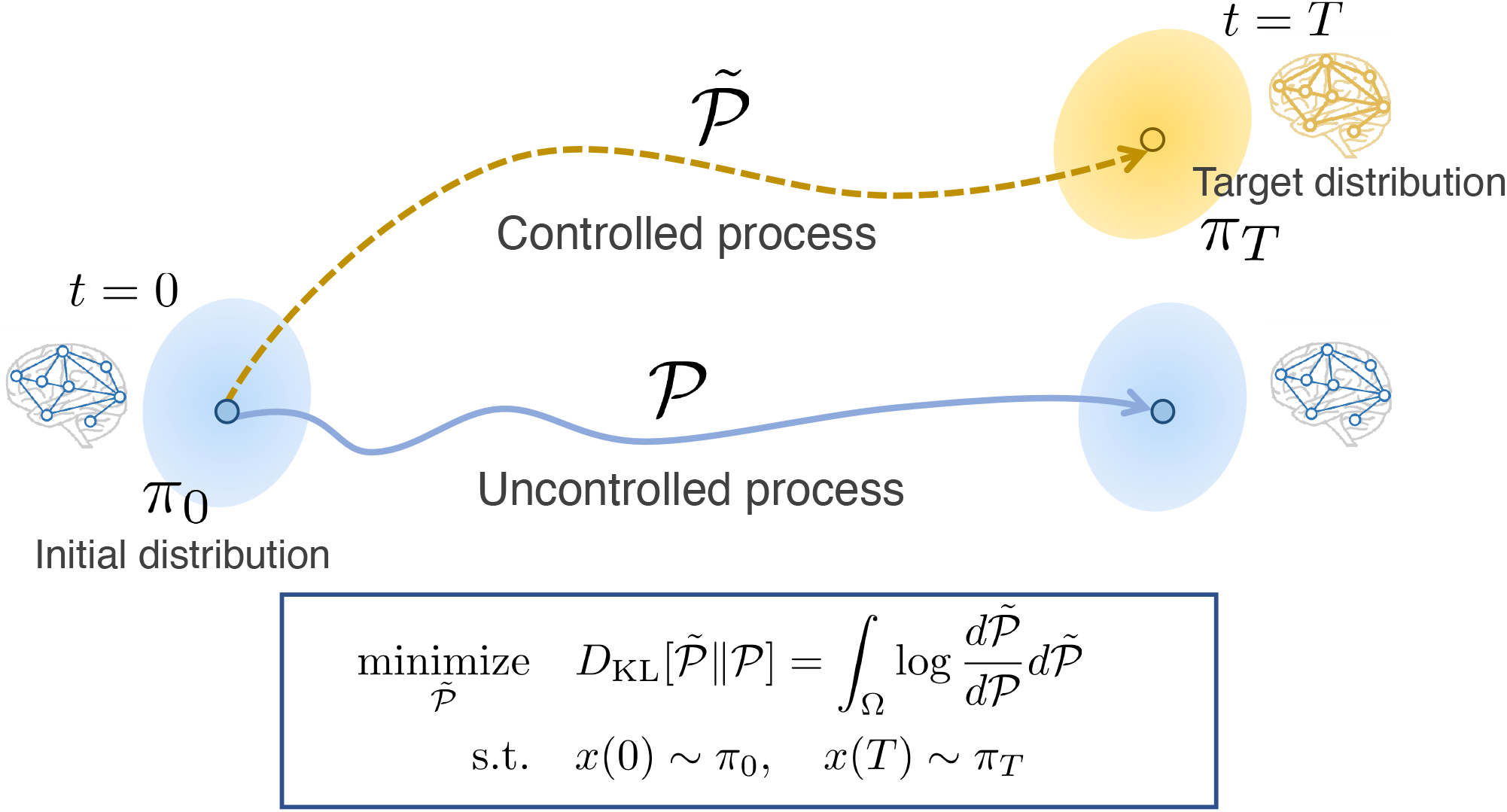
Schematic of our framework for quantification of control costs in the brain in linear stochastic systems. We model a brain state to follow a certain probability distribution *π*_0_ at time *t* = 0 (blue ellipse on the left). In uncontrolled dynamics, the brain state stays in the same distribution (blue trajectory & blue ellipse on the right). However, in a state transition, the brain dynamics changes so that it reaches a target distribution *π*_*T*_ at time *t* = *T* (gold ellipse). We call this altered trajectory the controlled process (gold trajectory). To evaluate how close the controlled process is to the uncontrolled one, we employ the KL divergence between the two processes as the cost function, marginalized with the initial and the target distributions (blue square). The distributions of the processes are defined on a path space, the space composed of ℝ ^*n*^-valued continuous functions defined on [0, *T*]. This type of KL optimization problem on a path space is referred to as the Schrödinger’s bridge problem.

In addition to determining the analytical expression, we also showed that we can decompose this expression into the cost for controlling the mean (referred to as *mean control cost*) and that for controlling the covariance (*covariance control cost*). We proved that the mean control cost corresponds to the control cost in the previous deterministic setting. The covariance control cost, on the other hand, is the cost of controlling the co-variation among the system, which has not been quantified in previous studies in neuroscience.

After formulating the theoretical framework, we then posed the following two questions. First, how important is it to take account of covariance in estimating control cost? Second, what brain areas are significant in controlling brain state transitions? To address these questions, we applied our new method to real neural data. We used whole-brain fMRI BOLD data of 352 healthy adults, recorded as part of the Human Connectome Project (HCP) [18]. As for the first question, we found that the influence of the covariance control cost was indeed not negligible. And as for the second, we discovered that the lower visual areas and the posterior cingulate cortex (PCC) play important roles in controlling state transitions, but in different ways: the PCC acts in controlling the covariance and the lower visual areas act in controlling the mean in addition to the covariance.

## 2 Materials and Methods

### 2.1 Theoretical Background

#### 2.1.1 Control Cost in Deterministic Systems

Before formulating the control costs in stochastic systems, we start with a conventional deterministic framework and a method for quantifying control cost under this deterministic setting. The dynamics of the brain is modeled with an *n*-dimensional state space model, where a brain state *x* ∈ ℝ^*n*^ at time *t* ∈ [0, *T*] (*T* > 0) consists of *n* scalar values that represent the magnitudes of brain activity. Then for each *t* ∈ [0, *T*], *x* (*t*) is assumed as a point in an *n*-dimensional Euclidean space. A state transition of the brain is described as a trajectory on which a point travels from one state to another. An uncontrolled transition is described using linear dynamics, such as

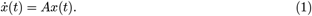

Here, *x* : [0, *T*] → ℝ^*n*^ is a vector that represents the magnitudes of activity of all nodes. *A* ∈ ℝ^*n*×*n*^ is a connectivity matrix whose elements represent connectivity weights for each pair of nodes.

We consider situations where a brain state switches from an initial state *x*(0) = *x*_0_ ∈ ℝ^*n*^ to a target state *x*(*T*) = *x*_*T*_ ∈ ℝ^*n*^ that is different from the state when following its uncontrolled dynamics (Eq. (1)). To realize such a transition, we assume that a control input *u*(*t*) is given. We can incorporate the control input into the dynamics as follows:

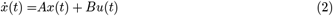

where *u* : [0, *T*] → ℝ^*p*^ is a control input and *B* ∈ ℝ^*n*×*m*^ is an input matrix that determines the nodes assigned with control inputs (*n, m* ∈ ℕ). We assume that this system is controllable (i.e., ∀*x*_0_, *x*_*T*_ there exists *u*(*t*) that enables the transition from *x*_0_ to *x*_*T*_). Here, we limit ourselves to cases where *B* = *I*_*n*_ (the *n*×*n* identity matrix), which implies that the input is given independently to all nodes. In other words, we consider the system with an input *v* : [0, *T*] → ℝ^*n*^,

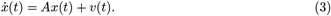

The control cost (also called control energy) 𝒥_cont_ is defined as the total amount of control input required to steer the system from *x*_0_ to the target *x*_*T*_ and is expressed as

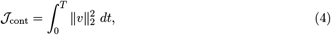

under

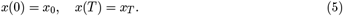

The minimum of this integral represents the input minimally needed to realize the state transition that satisfies the marginal conditions (Eq. (5)). It is written as

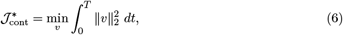

and referred to as the minimal control cost. The problem of minimizing the control cost is called the *optimal control problem* in control theory. Hereafter, we call the optimal control cost simply the “control cost” because we are only concerned with the minimal value in this study. We also refer to this cost as the deterministic cost, since this cost is defined on the deterministicm model (Eq. (3)). This minimum is known to exist, and the optimal value 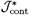 is readily solved (see [19])). This metric has been used as the control cost in previous studies in neuroscience [7, 12, 13, 14, 15, 20].

#### 2.1.2 Formulation of a Stochastic Linear Model

To take account of noise and fluctuation in brain activity [21], we make an extension for three characteristics in the control model of the brain: dynamics, transitions, and the control cost (Tab.1). In this section, we explain these characteristics in detail.

First, we model the brain dynamics through a stochastic process instead of conventional deterministic processes as described in Eq. (1) and Eq. (3). We then model the dynamics without control, or the *uncontrolled process* described as an ℝ^*n*^-valued Ornstein Uhlenbeck (OU) process *x*(*t*) (*t* ∈ [0, *T*]),

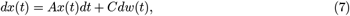

where each element of *x*(*t*) corresponds to the magnitudes of activities in a brain region. Here, *w*(*t*) is a standard *n*-dimensional normal Brownian motion, and *A* ∈ ℝ^*n*×*n*^, *C* ∈ ℝ^*n*×*n*^ (*n* ∈ ℕ). We assume *C* ^t^*C* to be nonsingular (^t^*C* denotes the matrix transpose of *C*), which is equivalent to rank(*C*) = *n*.

Second, to model the control of transitions, we consider these as from a probability distribution to another probability distribution, instead of as the point-to-point transitions that have been considered in previous studies [5, 9]. In the present study, we consider the control of the system between Gaussian distributions from time *t* = 0 to time *t* = *T*. We assume that the distribution of *x* at *t* = 0 follows a Gaussian distribution with mean *µ*_0_ ∈ ℝ^*n*^ and covariance Σ_0_ ∈ PSD(*n*) (set of *n*-by-*n* positive semi-definite matrices), denoted by *x*(0) ∼ 𝒩 (*µ*_0_, Σ_0_) (we call this the *initial distribution*). When the system follows Eq. (7), the distribution of *x*(*T*) is uniquely determined:

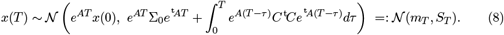

Specifically, if

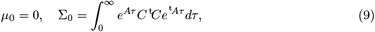

the process (Eq. (7)) stays in the same probability distribution in ∀*t* ≥ 0 (steady state distribution), and thus *µ*_0_ = *m*_*T*_ = 0 and Σ_0_ = *S*_*T*_. The existence of the steady state distribution is guaranteed if the real parts of all the eigenvalues of *A* are negative. To describe the control of state transitions, we consider altering the dynamics and the probability distribution at time *t* = *T* by giving a certain input. We consider steering the system from the initial distribution 𝒩 (*µ*_0_, Σ_0_) to a given distribution 𝒩 (*µ*_*T*_, Σ_*T*_) (*µ*_*T*_ ∈ ℝ^*n*^, Σ_*T*_ ∈ PSD(*n*)) at *t* = *T*, which is different from the final distribution 𝒩 (*m*_*T*_, *S*_*T*_) in the uncontrolled process. We call 𝒩 (*µ*_*T*_, Σ_*T*_) a *target distribution*, and a process that reaches 𝒩 (*µ*_*T*_, Σ_*T*_) at *t* = *T* a *controlled process*.

Third, for the control cost, we adopt the Kullback-Leibler (KL) divergence between the uncontrolled and controlled processes [6, 22, 23, 24]. A KL divergence is a metric that measures the closeness between two probability distributions. Thus, the control cost, defined by the KL divergence, measures the closeness between the uncontrolled and controlled processes. Note that this KL divergence is not the one between the two probability distributions of brain states at a particular time point, such as the KL divergence between the initial distribution 𝒩 (*µ*_0_, Σ_0_) and the target distribution 𝒩 (*µ*_*T*_, Σ_*T*_), but rather the divergence between the probability distributions of the entire paths in the uncontrolled and controlled process from time 0 to *T*.

In this study, we consider the optimal control problem based on the KL divergence. There are many possible controlled processes that take the system to a given target distribution. Among them, we consider the optimal controlled process, whose control cost is minimized. This process is the closest controlled process to the uncontrolled process in the KL sense. To express this mathematically, let us denote the probability distribution induced by the uncontrolled process by 𝒫 and that by the controlled process 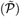 which are defined on an abstract space composed of a set of ℝ^*n*^-valued continuous functions on [0, *T*] (denoted by *S* = *C*([0, *T*], ℝ^*n*^)). The KL divergence between these paths is written as

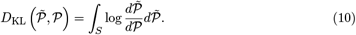

where 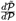 is a Radon-Nikodym derivative (we consider when 𝒫 and 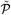 are absolutely continuous to each other). Then the probability distribution of the optimal controlled process we seek is the following *𝒬*:

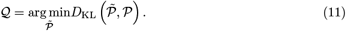

with the boundary conditions;

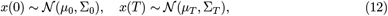

if the minimum exists. We call the minimum the *stochastic control cost*,

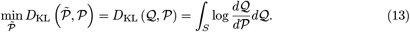

Again, the proposed framework is compared to the conventional deterministic framework in Table 1.

**Table 1:**
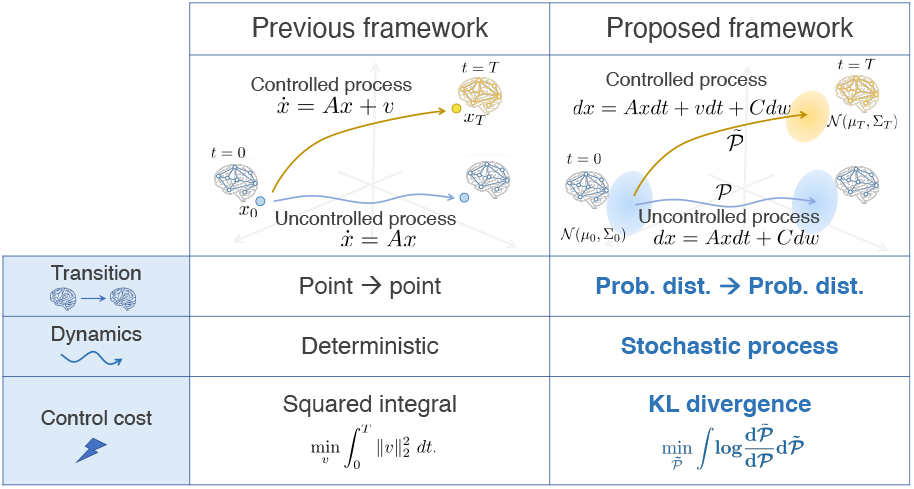
Comparison of deterministic and stochastic state transitions in a brain from the perspective of control theory. The **Previous framework** column describes the deterministic model of state transitions. The uncontrolled process (blue arrow) is expressed as a linear differential equation 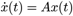 (*A* is a connectivity matrix of the brain network). A brain state transition is modeled as a transition from a point to another in an *n*-dimensional state space. Along with a control input *v*, the brain shifts its state to *x*_*T*_, another point in the state space, at time *T* (dynamics drawn in yellow). The cost function is often set as the time integral of the squared input. The **Proposed framework** column explains the stochastic model of brain state transitions. Here, the uncontrolled process (blue arrow) and a controlled process (gold arrow) are given as stochastic processes. A brain state transition is viewed as a shift from an *n*-dimensional probability distribution to another. As a control cost, the KL divergence between the two processes is examined.

|  | Previous framework | Proposed framework |
| --- | --- | --- |
|  | Controlled process<br>$\dot{x} = Ax + v$<br>Uncontrolled process<br>$\dot{x} = Ax$ | Controlled process<br>$dx = Axd t + vdt + Cdw$<br>$\tilde{\mathcal{P}}$<br>Uncontrolled process<br>$dx = Axd t + Cdw$<br>$\mathcal{P}$ |
| Transition | Point → point | **Prob. dist. → Prob. dist.** |
| Dynamics | Deterministic | **Stochastic process** |
| Control cost | Squared integral<br>$\min_v \int_0^T \ v\ _2^2 dt.$ | **KL divergence**<br>$\min_{\tilde{\mathcal{P}}} \int \log \frac{d\tilde{\mathcal{P}}}{d\mathcal{P}} d\tilde{\mathcal{P}}$ |

**Table 2:**
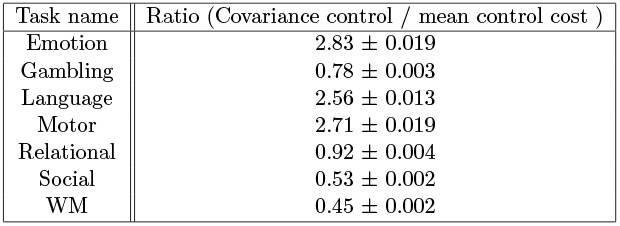
Ratios of covarariance control compared to the mean control cost in the transition from rest to each task. In each of the 100 bootstrap trials, the co-variance control cost is divided by the mean control cost, and the average and the standard deviations of the 100 trials are computed.

The optimization problem of minimizing the KL divergence between two stochastic processes (Eq. (11)) has been referred to as *Schrödinger*’*s bridge problem* [22, 23]. This problem was originally proposed by Erwin Schrödinger for finding the most probable path that moving particles take [25]. The Schrödinger bridge problem was subsequently found to be equivalent to an optimal control problem, and has been studied in the field of control theory [23, 24, 26]. To our knowledge, our recent study [6] is the first work in neuroscience to use the KL minimization problem in examining control cost in brain dynamics.

#### 2.1.3 Equivalence between KL cost and Quadratic Cost

Next, we observe that KL minimization boils down to an optimal control problem where the cost function is the expectation of the quadratic input with respect to the controlled process [24, 27].

It is known from previous studies [24, 26] that to consider the minimal KL divergence 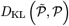 under the boundary conditions (Eq. (12)), one has only to search for dynamics that can be described as the next form

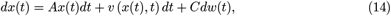

where *v* : ℝ^*n*^ × [0, *T*] → ℝ^*n*^ is a control input. Moreover, the KL divergence and the next quadratic cost become equal:

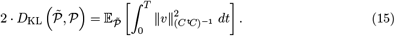

where 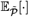 represents the expectation on a probability law 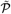 and 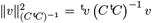.Therefore, to obtain the minimal KL divergence, we need to search for the input *v* that minimizes the expectation of the quadratic form given in the right hand side of Eq. (15). In other words, the KL minimization problem boils down to an optimal control problem whose cost function is the right hand side of Eq. (15). As defined in the previous section, we define the minimum of Eq. (15) as the *stochastic control cost*.

The discussion in this section is stated in more rigorous form in [19].

### 2.2 On fMRI Data

The 3T functional magnetic resonance imaging (fMRI) data of 990 subjects were obtained from the Washington University-Minnesota Consortium Human Connectome Project (HCP) [18]. To remove the effects of outliers, we picked data of 352 subjects according to criteria suggested in [28].

### 2.3 Data Preprocessing

Minimally preprocessed fMRI data were used for the resting state and seven cognitive task states (emotion, gambling, language, motor, relational, social, and working memory). Denoising was performed by estimating nuisance regressors and subtracting them from the signal at every vertex [29]. For this, 36 nuisance regressors and spike regressors were used following a previous study [29], consisting of (1-6) six motion parameters, (7) a white matter time series, (8) a cerebrospinal fluid time series, (9) a global signal time series, (10-18) temporal derivatives of (1-9), and (19-36) quadratic terms for (1-18). The spike regressors were computed with 1.5 mm movement as a spike identification threshold. After regressing these nuisance time courses, a band-pass filter (0.01-0.69 Hz) was applied, whose upper filter bound corresponds to the Nyquist frequency of the time series. A parcellation process proposed in [30] was used to divide the cortex into 100 brain regions, which reduced the complexity of the following analysis.

### 2.4 Estimation of Parameters Characterizing Brain States

We next estimated parameters that characterize resting and task brain states. To apply the framework explained in Section 2.1.2, we estimated the next parameters: the mean and the covariance matrix of the initial distribution (*µ*_0_ and Σ_0_), those of the target distribution (*µ*_*T*_ and Σ_*T*_), and the matrices in the uncontrolled dynamics (the drift matrix *A* ∈ ℝ^*n*×*n*^ and the product of the diffusion matrix *S*_*C*_ := *C* ^t^*C*). As for the diffusion matrix, we estimated *S*_*C*_ = *C* ^t^*C*∈ ℝ^*n*×*n*^ rather than *C* itself, as *S*_*C*_ is sufficient for cost computation. The detailed methods to estimate the parameters of the initial and the target distributions are covered in 2.4.1, and those to estimate *A* and *S*_*C*_ in 2.4.2. For statistical robustness, a bootstrapping method was adopted, where the estimation described below was done using data of 100 randomly chosen subjects out of 352 overall subjects, and repeated 100 times independently. The data of the 100 subjects were concatenated to obtain a single time series that is long enough for statistically reliable estimation.

Before estimating the parameters using the concatenated data across different subjects, we need to normalize the time series data. This normalization is necessary for the following reason. Basically, the absolute values of fMRI BOLD signals are meaningless by themselves, due to the bias caused by the water content or the basic blood flow in the brain tissues [31]. Because of this, one cannot directly compare values of BOLD signals in different subjects. Thus, we cannot concatenate data of different subjects using the preprocessed fMRI BOLD signals as they are. To concatenate BOLD signals across subjects, we need some kind of scaling, or normalization, of the data.

We performed the scaling based on the assumption that the sum of the time series variances in the whole ROIs are constant when a subject is not engaged in a cognitive task. First, for each subject, we computed the trace values of the empirical covariance matrices of the resting state and the task-free moments in each task. Then, we divided the whole time series data of the rest and each task by the square root of the trace value of the rest and by that of each task’s task-free moment, respectively. This division makes the magnitude of the sample covariance matrix (i.e., the trace of the sample covariance matrix) of the rest and the task-free moments to be one in all subjects and tasks. After this normalization, we concatenated the time series data of the whole resting state and the task-free moments of each task state of all 100 subjects. Using the concatenated time series data, we estimated the parameters as follows.

#### 2.4.1 Distribution of the Resting State and Task states

To estimate the resting state distribution, the empirical mean and covariance of the whole time series were used to represent the probability distribution. The empirical mean of the resting state was almost a 100-dimensional zero vector in each subject due to a zero-mean adjustment in preprocessing. To extract the mean of the activity during a task, we did not use the empirical mean and covariance as we did in estimating the resting state distribution. This is because unlike the resting state data, task time series data are composed of task-performing and task-free moments. And one needs to know to what extent the activity in the task-performing moments is different from that in the task-free moments. We assumed that in the task-free moments, the average magnitude of activity is the same as the average of the resting state, i.e., approximately zero. Accordingly, to obtain the time series mean of the activity during the task state, the time series mean during the task-free moments was subtracted from that in the task-performing moments. The covariance matrix was calculated as the empirical covariance matrix during the task-performing moments.

#### 2.4.2 Estimation of Parameters of the Resting State Dynamics

In this section, we explain how we estimated the matrices *A* and *S*_*C*_ = *C* ^t^*C* that determine the uncontrolled dynamics (Eq. (7)). As we mentioned in 3.2, we regarded the uncontrolled dynamics as the resting state dynamics. Thus we need to fit the resting state time series to the dynamics equation

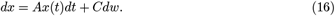

When sampled at some interval of Δ*t* > 0, this process is equivalent to the next VAR(1) process,

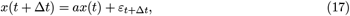

where *a* = *e*^*A*Δ*t*^ (matrix exponential). We first estimated the drift term (denoted by *â*) using the least absolute shrinkage and selection operator (LASSO) regression. A transformation

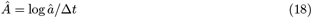

followed to obtain the estimated drift coefficient *A* in the dynamics.

As for *S*_*C*_ = *C* ^t^*C*, one can directly infer this from the covariance of 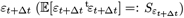 in equation (17) using the relationship

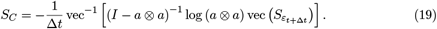

where vec is the vectorization operator and vec^−1^ its inverse, and log the matrix logarithm. To obtain the estimation of *S*_*C*_ (denoted by *Ŝ*_*C*_), we substituted above the estimated *â* and the empirical time series covariance 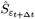.

### 2.5 Calculation of Entropy of an Input Map

The entropy *S* of an input map *I* = ^t^(*I*_1_, · · ·, *I*_100_) ∈ ℝ_+_^100^ quantifies how dispersed a vector is and takes the maximum if each element *I*_*l*_ (*l* = 1, · · ·, 100) takes the same value. *S* is calculated by

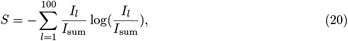

where 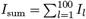. Note that each element *I*_*l*_ (*l* = 1, · · ·, 100) is positive.

### 2.6 Code Accessibility

The codes for computing the stochastic control cost and reproducing the figures on this paper are available at https://github.com/oizumi-lab/SB_toolbox.

## 3 Results

### 3.1 Theoretical Results

In this section, we show theoretical results obtained from our framework proposed in the previous section. This framework based on the KL minimization was first introduced in neuroscience in our previous study [6]. While the previous study considered a finite-state space, discrete-time stochastic process, we considered a linear continuous-time system in this study. As we have explained in the previous section, there have been many theoretical studies of the linear stochastic system. In addition to what has previously been acknowledged, we newly obtained the following analytical results:

1. We derived the analytical solution of the control cost. We found that the analytical solution of the stochastic cost can be disintegrated into two portions, the cost of driving the mean (mean control cost) and that for the covariance (covariance control cost).
2. The mean control cost turns out to correspond to the conventional deterministic control cost in specific occasions. This clarifies the correspondence between the deterministic and the stochastic costs. The covariance control cost has not been quantified in previous applications in neuroscience.
3. By investigating the control input assigned to each node, we can compute the amount of total input given at one node. This node-level input can also be decomposed into two parts, the input necessary for controlling the mean and that necessary for controlling the covariance.

#### 3.1.1 Analytical Solution of the Stochastic Control Cost

We start by showing the analytical solution of the optimal value of Eq. (15), which is described as follows:

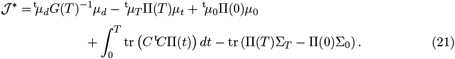

In the equation, *G* : [0, *T*] → ℝ^*n*×*n*^, Π : [0, *T*] → ℝ^*n*×*n*^, Ψ : [0, *T*] × [0, *T*] → ℝ^*n*×*n*^, and *µ*_*d*_ := *µ*_*T*_ − Ψ(*T*, 0)*µ*_0_. See [19] for the definitions and detailed derivations. Note that Π(*t*), *G*(*t*), and Ψ(*t, s*) depend on *A, C*, Σ_0_ and Σ_*T*_. This gives the analytical expression of the stochastic control cost.

#### 3.1.2 Decomposition of the Stochastic Cost

We next demonstrate that Eq. (21) can be decomposed into two parts: the cost needed to steer the mean (called *mean control cost*) and the cost needed to steer the covariance (called *covariance control cost*). More specifically, when we decompose this as

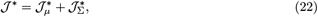

where

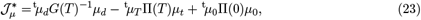

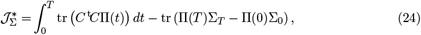

we show that we can consider 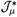 as the cost needed steer the mean (mean control cost), and 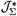 as the cost needed to steer the covariance (covariance control cost). This decomposition is shown schematically in Fig. 2.

**Figure 2:**
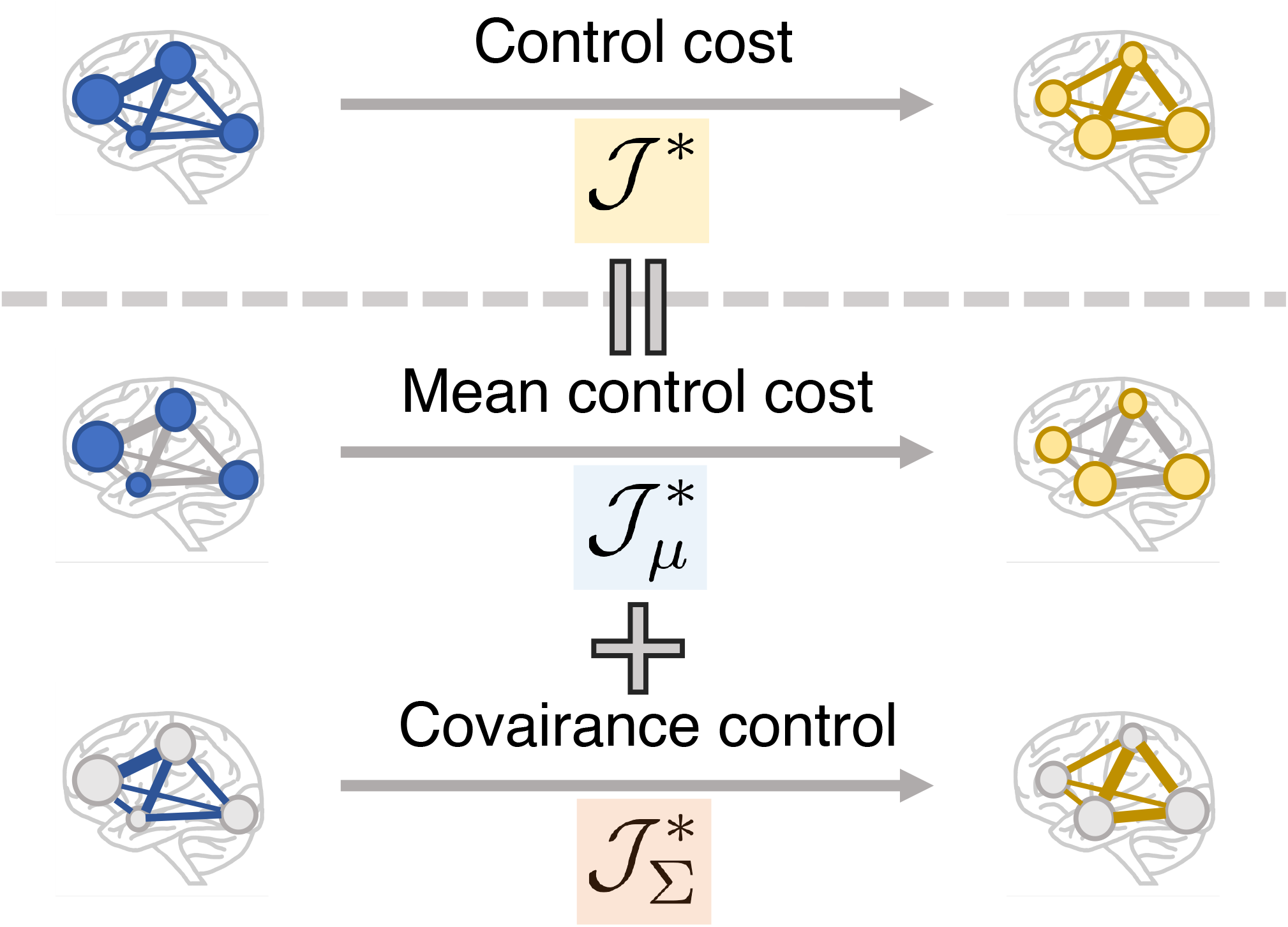
Schematic illustration of the decomposition of stochastic control cost into the mean and covariance control. The stochastic control cost from one state to another is described as a sum of the cost to control the mean and covariance from the initial state to the target state. The radius of the circle represents the magnitude of the mean, while the thickness of an edge represents the covariance value between two nodes.

We found that the mean control cost 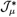 depends only on the marginal means (*µ*_0_ and *µ*_*T*_), while the covariance control cost 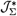 depends only on the marginal covariances (Σ_0_ and Σ_*T*_). To explain these dependencies, we start by examining 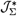. We can see that Eq. (24) depends only on the marginal covariances and is independent of the marginal means, since Π(*t*) relies only on Σ_0_ and Σ_*T*_ (in addition to *A* and *C*). We can then interpret these terms to represent the cost of navigating the covariance from Σ_0_ to Σ_*T*_. On the other hand, 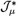 (Eq. (23)) seems to depend on both the marginal means and marginal covariances, since *G* and Π are dependent on Σ_0_ and Σ_*T*_. Surprisingly, however, we found that Eq. (23) actually does *not* depend on the marginal covariances, but rather only on the means. In fact, 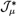 is further rewritten as

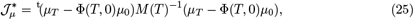

where Φ is a state transition matrix defined as Φ(*t, s*) = *e*^*A*(*t*−*s*)^, and *M* (*T*) is a Gramian matrix in the linear stochastic system,

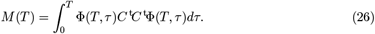

See [19] for the proof of this transformation (Eq. (25)). Thus, Eq. (23) can be seen as the cost for controlling the mean from *µ*_0_ to *µ*_*T*_.

Furthermore, the formulation given in Eq. (25) indicates a correspondence between the stochastic control cost and the deterministic control cost [19]. We can easily see that if the input matrix *B* is identical to the diffusion matrix *C*, the mean control cost (Eq. (25)) is equal to the deterministic control cost. This correspondence between the mean control cost and the deterministic control cost indicates that the stochastic control cost, defined as the KL divergence, can be considered an extension of the conventional deterministic control cost.

Another merit of the expression of the stochastic cost (Eq. (25)) using the the Gramian matrix (Eq. (25)) is that one can obtain the directions in which controlling the mean is easy or difficult. In the deterministic control system, it is classically known that the eigenvectors of the control Gramian matrix (see [19]) can be interpreted as the direction in which the control is easy/difficult [32, 33]. The same logic can be applied to the mean control cost that shares the similar formulation to the Gramian in the deterministic system: the eigenvectors of Eq. (26) specify the directions in which the system is good at in controlling its mean.

The covariance control cost, on the other hand, is the cost of shifting the covariance of the probability distribution. This cost has been ignored in previous neuroscience studies [5, 7, 9, 34, 13, 15]. We will cover the neuroscientific implication of the covariance control cost, which can be rephrased as the cost for controlling functional connectivity, in the Discussion. As the formulation Eq. (24) is somewhat complicated, an intuitive interpretation of the covariance control cost is not available at this moment.

The significance of this decomposition is that it enables us to separately quantify the influence of taking into account the covariance of brain activities in the control cost (Fig. 2). As we saw, the mean control cost depends only on the marginal means and is shown to correspond to the deterministic control cost under a certain setting. On the other hand, the covariance control cost depends only on the marginal covariances. Thus, the effect of taking account of the covariance of brain activities is reflected only in the covariance control cost.

#### 3.1.3 Input Given at Each Node

Lastly, we evaluate the contribution of each node to control transitions by the total amount of inputs given at each node in the brain network.

The amount of inputs provided at each node can be calculated as follows. Again, the optimal dynamics is described as

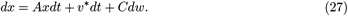

We let 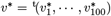 where 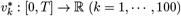. The expectation of the total input given to the *k*’th node (denoted by ℐ(*k*), *k* = 1, · · ·, 100) is then given as

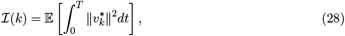

which is readily transformed into

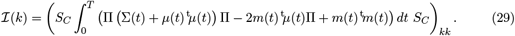

The subscript (·)_*ij*_ denotes the (*i, j*)-th entry of a matrix, and *m* : [0, *T*] → ℝ^*n*^ is a function for arranging the transient mean. See [19] for the derivation, and *m* : [0, *T*] → ℝ is a certain function defined in [19]. This value can be decomposed in a similar way to the stochastic control cost (Eq. (22)),

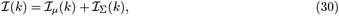

where

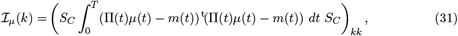

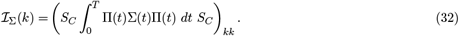

As *m*(*t*) is the function for shifting the mean of the brain dynamics, the first term ℐ_*µ*_(*k*) can be interpreted as the cost needed to control the mean, which we call the *mean input* at the *k*’th ROI. The latter ℐ_Σ_(*k*) is the cost of controlling the covariance, which we refer to the *covariance input* at the *k*’th ROI.

### 3.2 Results on Application to fMRI Data

So far we have formulated a method to quantify the stochastic control cost and contribution of each brain region on state transitions. With this method, we then address the following two problems:

- How important is it to take account of the covariance of brain state probability distributions?
- What brain areas are significant in optimally controlling the brain state transitions?

To address the first question, i.e. to assess the influence of covariance, one can compare the covariance control cost 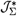 to the mean control cost 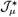. We have seen that the mean control cost in the stochastic model corresponds to the optimal control cost in the deterministic model (Section 3.1.2), where we consider the control from one point to another point. So imagine, for instance, a case where the mean control cost is far greater than the covariance control cost and hence accounts for the most part of the overall stochastic cost. We might then need to consider only the point-to-point control and might not have to investigate distribution-to-distribution control as we did in the stochastic model. To evaluate the necessity of incorporating probability distributions in our model, we first need to examine to what extent the covariance control cost accounts for the stochastic cost.

To address the second question regarding important brain regions in controlling brain state transitions, we computed control input *v* for each region as defined in Section 3.1.3. By decomposing the control input of each region into the mean input (ℐ_*µ*_(*k*)) and covariance input (ℐ_Σ_(*k*)), we can identify regions that are important for shifting the mean and covariance of brain activity. This decomposition enables us to identify not only the brain regions that need to be activated but also those regions that are crucial to the reconfiguration of co-variation of whole brain activity.

To address these questions, we used whole-brain functional magnetic resonance imaging (fMRI) data recorded as part of the Human Connectome Project (HCP) [35]. These data consist of recordings of 990 healthy adults. We used a subset of data composed of 352 subjects based on a criteria proposed in [28]. For each subject, the dataset contains blood-oxygen-level-dependent (BOLD) signals, including a pair of scans at rest and scans under seven different cognitive tasks: emotion, gambling, motor, language, relational, social, and working memory. After a standard preprocessing procedure including denoising with nuisance regressors, the whole voxels were partitioned into 100 regions of interest (ROIs) (Fig. 3-a & b) following the method proposed in [30].

**Figure 3:**
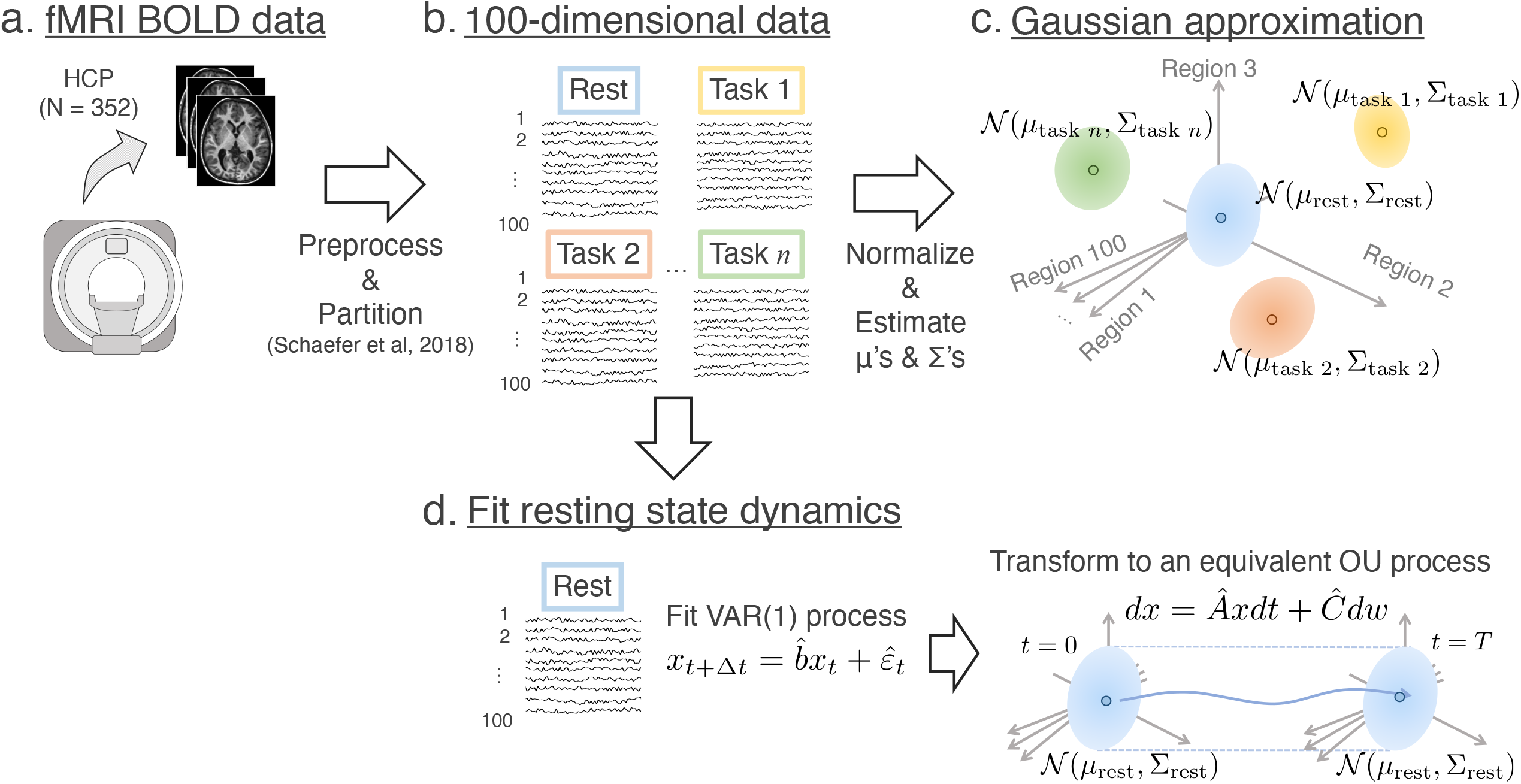
Data acquisition techniques. **a**. The fMRI BOLD data in HCP underwent minimal preprocessing and were then divided into 100 regions of interest (ROI). **b**. This gives us 100-dimensional time series data in our hands. **c**. The data at rest and under each task state were approximated as a Gaussian distribution in the 100-dimensional space whose individual coordinates represent the magnitude of activity of an ROI.

After preprocessing and parcellation, we applied our framework to the fMRI data. Here, we assumed that the uncontrolled process is the resting state dynamics of the brain and the controlled dynamics is the transition from the probability distribution of the resting state to that of a task state. Under this assumption, we computed the control cost from the resting state probability distribution to a task distribution. To perform this computation, we need to infer two classes of probability distributions: i) the probability distributions that characterize the resting state dynamics, and ii) the task state probability distribution. For i), we employed a regression using the sparse vector autoregressive (VAR) model (Fig. 3-d) on the resting state BOLD data. We assumed the resting state dynamics to be the steady state dynamics and computed the steady state probability distribution (𝒩 (*µ*_0_, Σ_0_)) and the transition probability distribution (characterized by the matrices *A* and *C*). For ii), the mean (*µ*_*T*_) and covariance (Σ_*T*_) are inferred using the sample mean and covariance in the BOLD signals after normalization (Fig. 3-c).

As for the target time constant, we used various *T* values ranging from 0.1 to 6.0 seconds, and observed that results did not qualitatively change [19]. Thus, below, we show the results with *T* = 1.0 seconds as representative.

To estimate the probability distributions and the stochastic costs, a bootstrapping method was adopted for statistical robustness. The estimation was carried out with 100 randomly chosen subjects out of 352 subjects and was repeated 100 times independently. See the Materials and Methods for further details.

#### 3.2.1 Mean and Covariance Control Cost in the Stochastic Model

Using the estimated probability distributions and the matrices *A* and *C*, we computed the stochastic control cost from rest to the seven tasks. For each task, we separately computed the mean control cost 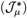, the covariance control cost 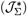, and their ratio 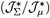.

First, to assess the contribution of the covariance control cost compared to the mean control cost, we show the ratio in Tab. 2. We found that covariance control costs are as large as, or slightly larger than, the mean control costs. The ratios range from 0.4 to 2.8 depending on the task, with an average value of about 1.5. This result indicates that the contribution of covariance control cost to the overall control cost is as important as that of mean control cost. We also found that these ratios of covariance control cost to mean control cost take various values depending on the specific task. For example, the language task needs a larger covariance control costs than the mean, while the opposite is the case for WM.

To identify the influence of the covariance control cost, we then examined if incorporating the covariance control cost changes the ordering of the seven tasks by the control costs. We found that incorporating covariance control cost changes the ordering of the seven tasks by the control costs. We show the mean, the covariance, and the stochastic control cost (= mean control cost + covariance control cost) of the seven tasks in ascending order (Fig. 4). The left panel (blue bars) shows the mean control cost values, the middle panel (orange bars) the covariance control cost, and the right panel (yellow bars) the total stochastic control costs. We can see that the order changes when we consider covariance control cost in addition to mean control cost. For example, the mean control cost for the gambling task is larger than that of the motor task; however, when we take account of covariance control cost, this relationship reverses. Thus, we see that the covariance control cost changes the result when estimating the magnitude relationships of control costs.

**Figure 4:**
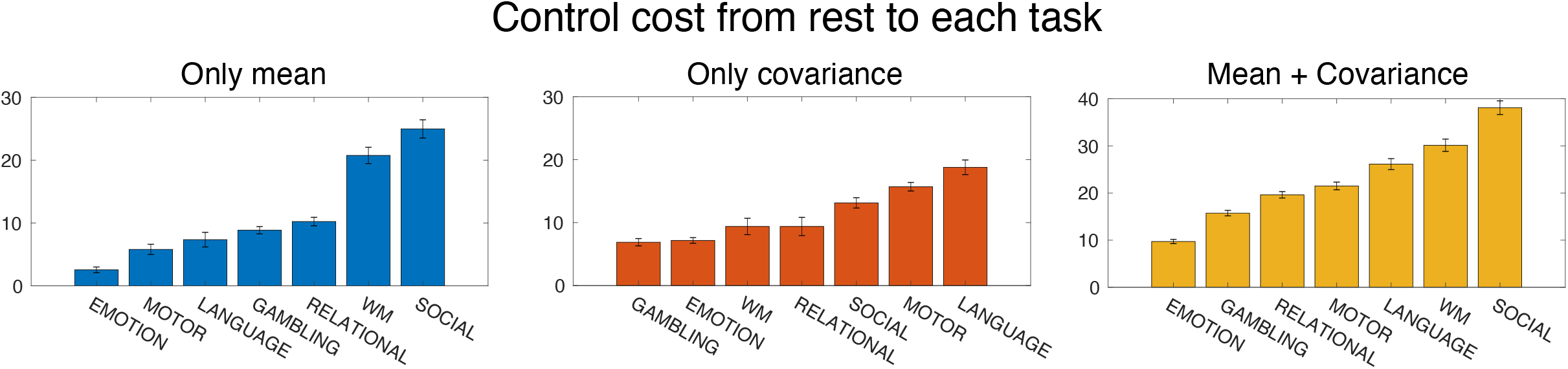
Stochastic control cost from the resting state to task states. Left panel, mean control cost values 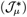; middle, covariance control values 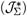; and right, total control values 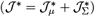 in ascending order.

#### 3.2.2 Control Inputs at Brain Regions

To identify which areas are important in optimally controlling brain state transitions, we next computed the amount of inputs given to each brain region (see 3.1.3). We computed the mean input *I*_*µ*_(*k*) (Eq. (31)) and covariance input ℐ_Σ_(*k*) (Eq. (32)) at each ROI, obtaining a pair of 100-dimensional vectors ((ℐ_*µ*_(1), · · ·, ℐ_*µ*_(100)), (ℐ_Σ_(1), · · ·ℐ_Σ_(100))) and their sum ((ℐ (1), · · ·, ℐ (100))). We can think of these vectors as brain maps of the amount of input required to alter the mean and covariance, and the total inputs. We call these maps the *mean input map, covariance input map*, and *total input map*. We computed the inputs and obtained these three maps for each of the seven tasks.

We found that the mean and covariance input maps showed quantitatively different patterns in each task. The mean inputs are typically located in a small number of regions, while the covariance input maps are more widely distributed throughout the brain [19]. We quantitatively evaluated the difference in distributions of the input maps by computing the entropy of the input maps over 100 ROIs for each task. The entropy *S* of a vector quantifies how dispersed the elements of the vector are; if the inputs are assigned to widely distributed regions, the entropy takes a large value. See the Materials and Methods section for a detailed description of how the entropy was calculated. As shown in Fig. 5, the entropy of the covariance input maps is larger than that of the mean input maps in all seven tasks. This result suggests that the covariance input maps are more dispersed over the whole brain regions.

**Figure 5:**
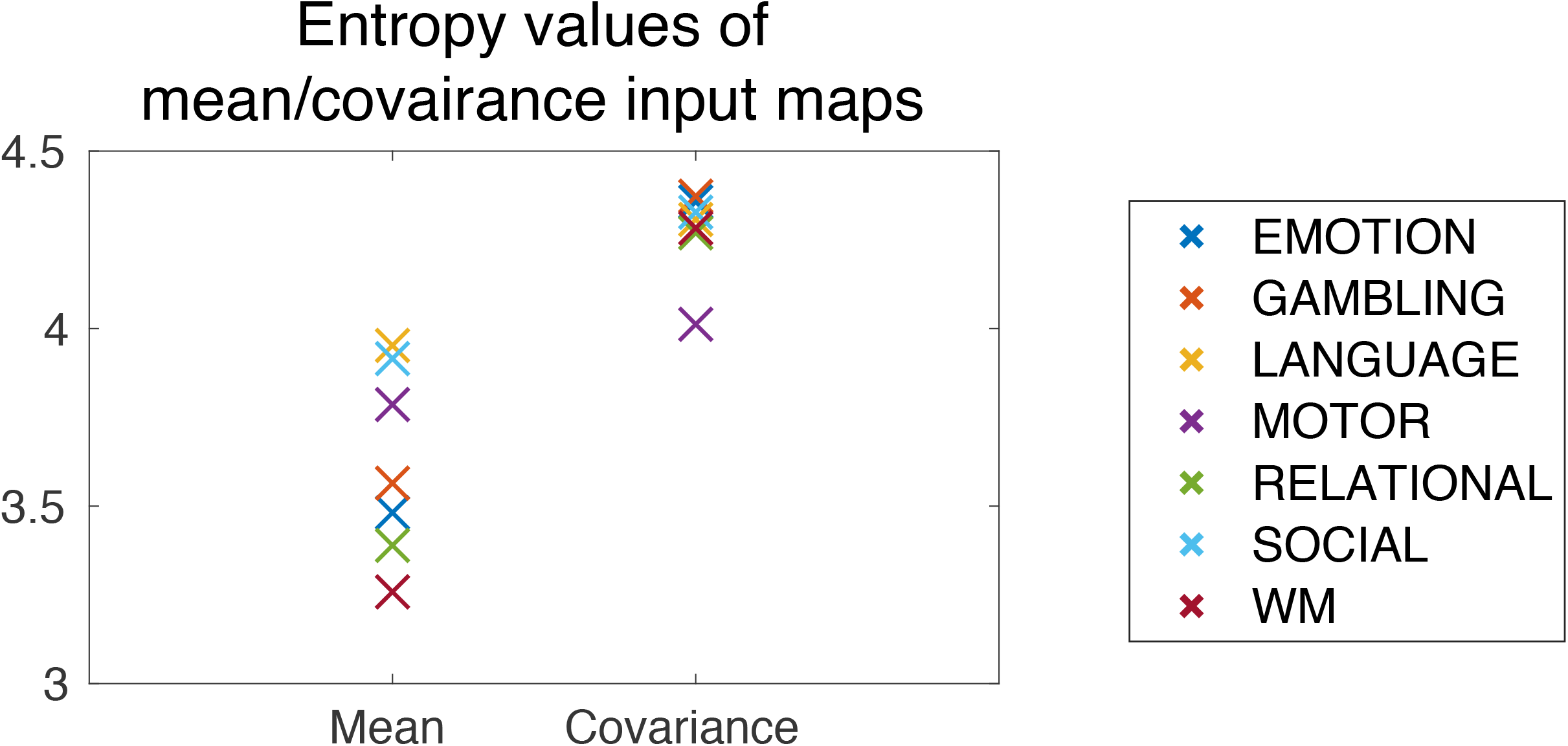
Entropy values of the mean and covariance input maps for each task. The values were averaged over 100 bootstrap trials.

We then examined the relationships between the two types of control inputs and the changes in BOLD signal magnitudes when transitioning to a task. Intuitively, we may expect that larger inputs are required for brain regions that are activated or deactivated in the task. To study this relationship, we computed the correlation coefficient of the the brain activity change and the mean/covariance input maps. We subtracted BOLD signals of preparation periods from those of task periods, and then computed the t-values of the differences. We defined the absolute value of the t-values as the brain activity change. We found that the mean input maps have high correlation coefficients in all the tasks, whereas the covariance input maps have lower values (Fig. 6). This result indicates that the mean input maps are highly correlated to the changes in BOLD activation level, while the covariance input maps are less related.

**Figure 6:**
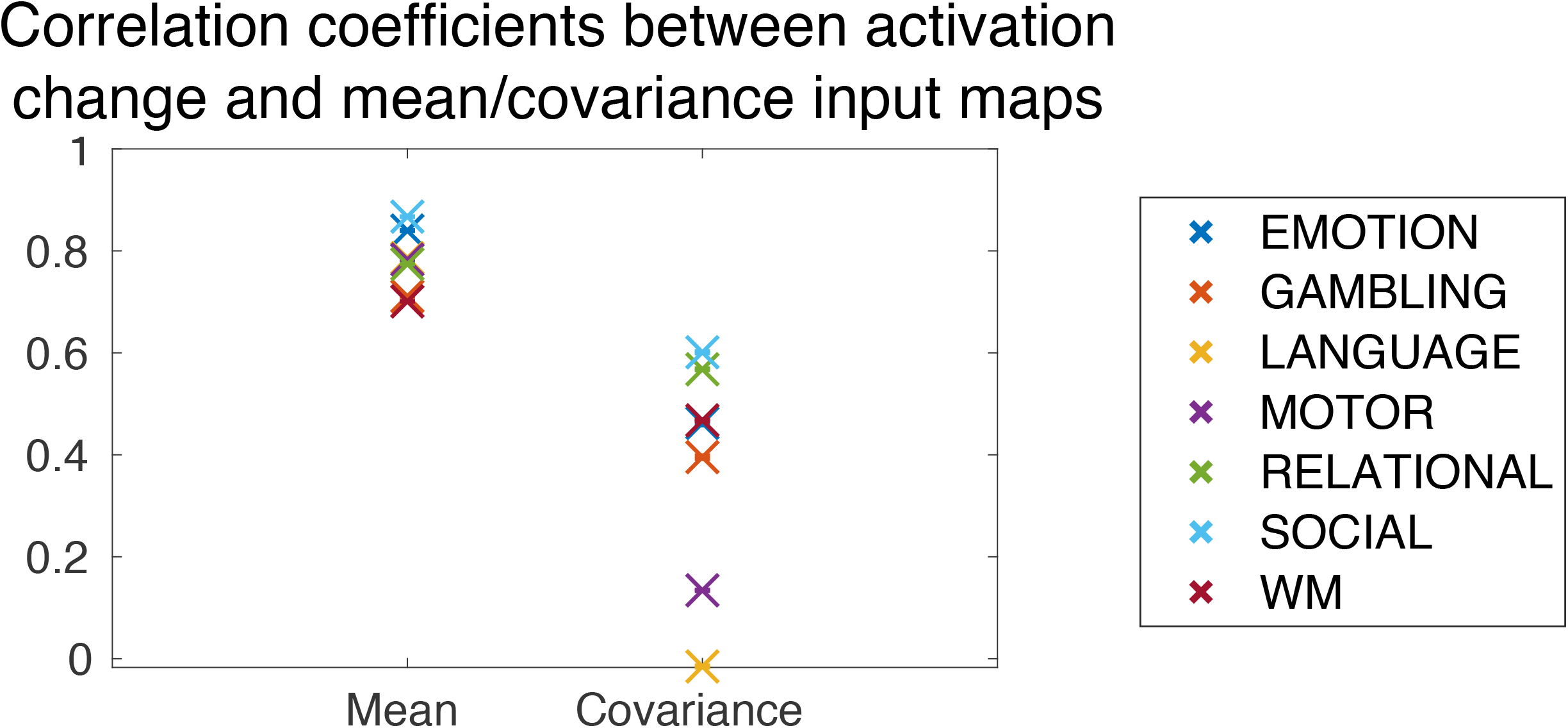
Correlation coefficients between the absolute values of t-values of activation and the input maps. The values were averaged over 100 bootstrap trials.

To identify the brain regions that are commonly important for control of the seven cognitive tasks, we then computed the average of the mean, the covariance, and the total input maps for all seven tasks, as shown in Fig. 7-a. To calculate the contribution of each task in a fair manner, for each task, we normalized the mean (ℐ_*µ*_(*k*)), the covariance (ℐ_Σ_(*k*)), and the total input map *I*(*k*), *k* = 1, · · ·, 100 by dividing by the sum of the total input to the whole ROIs 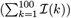, and then took the average of the seven tasks. To visually inspect particularly important ROIs, we created Fig. 7-b, which shows ROIs having the largest inputs and explaining 30% of the sum of each input to the whole ROIs 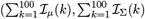, or 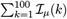. We can see that a large amount of the mean inputs is in the visual areas (Fig. 7-a, “mean”). In contrast, the covariance inputs are slightly more distributed over the whole brain. Specifically, a large portion of inputs is concentrated in the visual area, orbitofrontal area, and the posterior cingulate cortex (PCC) (Fig. 7-a “covariance”).

**Figure 7:**
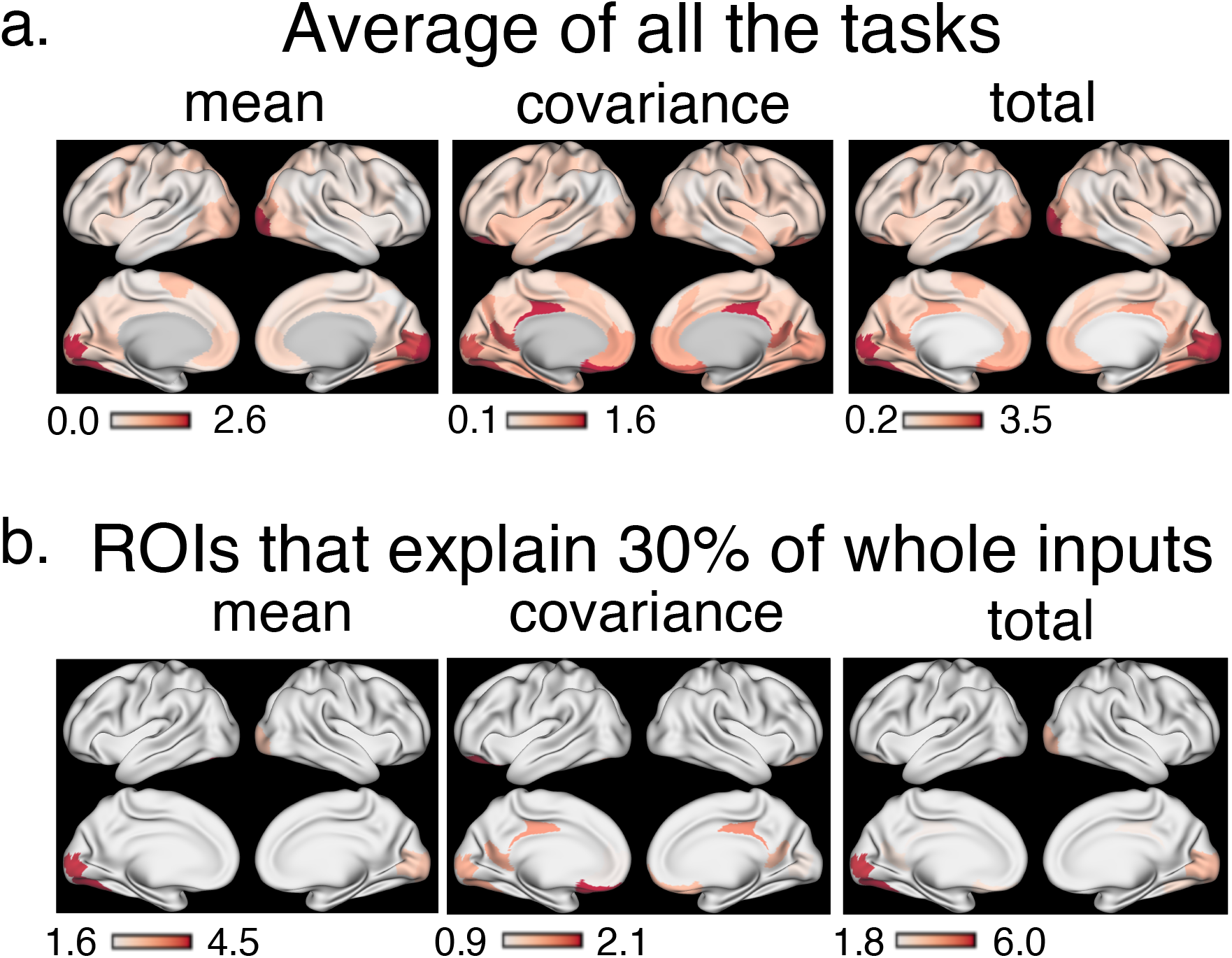
Mean, covariance, and total input maps averaged over the seven tasks. **a**. Average of the mean, covariance, and total inputs for each 100 ROIs over seven tasks in the HCP dataset. The numerical values indicate percentages compared to the sum of the total input to the whole ROIs. **b**. The ROIs that account for 30% of the whole mean, covariance, and total inputs shown in Fig. 7-a. The numerical values indicate the percentages compared to the sum of the mean, covariance, and total input to the whole ROIs, respectively.

To validate these results on the brain regions which are important in regulating brain state transitions, we counted how many times each ROI appears in the top 10 ROIs with the largest control inputs. Although we identified commonly important regions in controlling transitions by computing the average control inputs of all tasks, the possibility exists that the control input to a certain area is extremely large for one task only and not for the others. To rule out such a possibility and to find those regions which are commonly significant for the seven tasks, we selected the top 10 significant brain regions for each task and counted how many times each region is ranked in the top 10. We performed this analysis for the mean, the covariance, and the total inputs.

This additional analysis further supported our findings that the PCC and lower visual areas are generally important in control transition to task states. The results of the additional analysis are shown in Fig. 8. The color of an ROI signifies the number of tasks for which the ROI is ranked in the top 10 (thus the numbers take integer values from zero to seven). To facilitate visualization, we only colored ROIs that appeared in the top 10 more than three times. For mean input, the lower visual areas are ranked in the top 10 in the majority of the tasks. For the covariance input, the posterior cingulate cortex (PCC) is ranked in the top 10 most frequently, followed by the lower visual areas. For the total input, the lower visual areas and the PCC were ranked most important. Taking these findings together, we conclude that the PCC and the lower visual areas are the most significant regions for transitioning to task states. While the lower visual areas contribute to both the mean and covariance shift, the PCC contributes to shifting covariance only.

**Figure 8:**
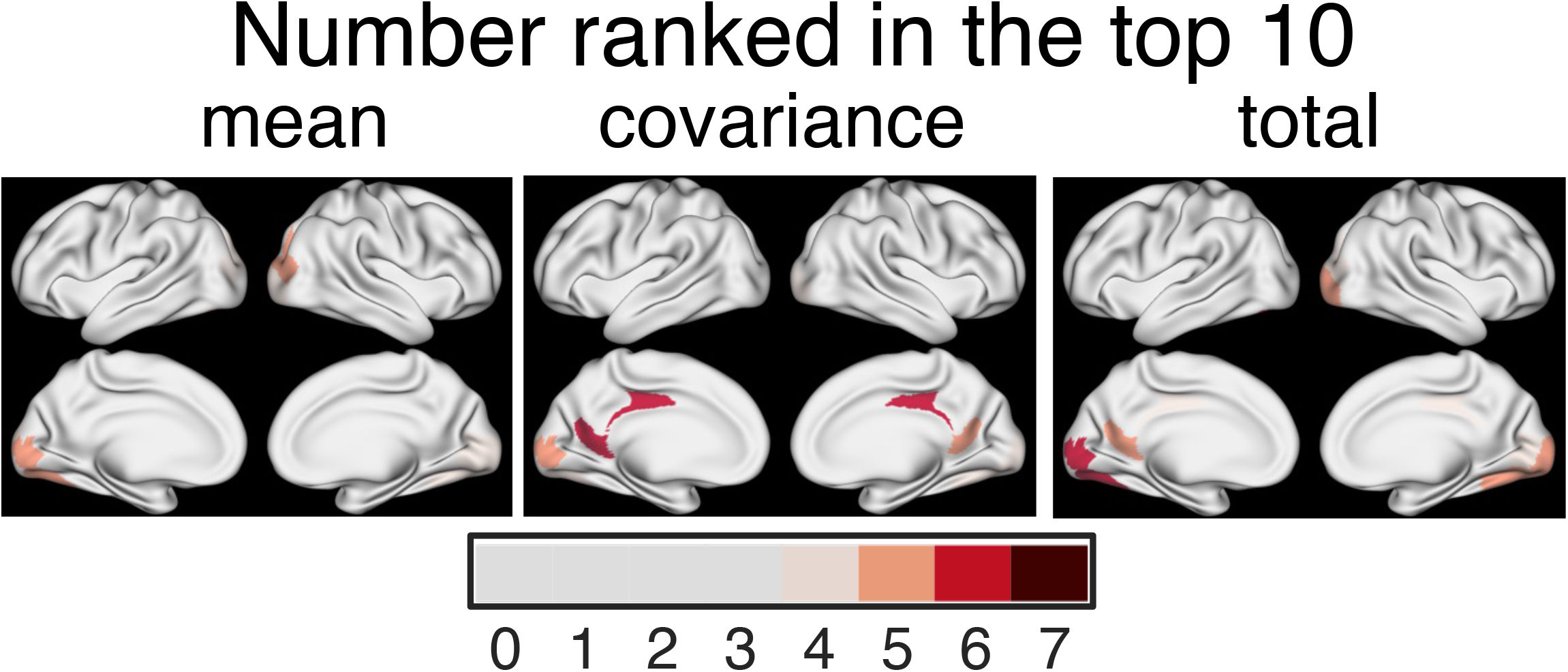
Number of times an ROI was ranked in the top ten brain regions that contributed most from rest to the seven tasks. From left to right, each panel shows the areas that contribute to the mean, covariance and total control, respectively.

## 4 Discussion

In this study, we propose a novel framework based on a linear stochastic dynamics to quantify control costs between brain states. We define the minimal KL divergence between the uncontrolled and the optimally controlled paths (Eq. (10)) as the stochastic control cost. There are three theoretical contributions in this study. i) We give a solution for the stochastic control cost (Eq. (21)). ii) We show that the cost can be decomposed into two parts: the mean control and covariance control costs (Eq. (22) and Fig. 2). iii) The proposed framework allows us to measure the inputs given in each ROI, thanks to the explicit modeling of the input term (Eq. (28) & Eq. (29)). The input in each ROI can be further decomposed into inputs necessary for regulating the mean brain activity and the whole-brain functional connectivity (Eq. (30)).

We applied our new method to fMRI BOLD data in the HCP dataset. We first found that the influence of incorporating covariance in control costs is not negligible. We further discovered that for regulating the mean, the lower visual areas turned out to be the most significant, whereas for the covariance, the PCC and the lower visual were the most significant.

### 4.1 Stochasticity of Control Cost

One contribution of the present framework is incorporating stochasticity in quantifying the control cost, which has not been adequately addressed in previous studies in neuroscience [5, 7, 9, 34, 13]. Our framework incorporates stochasticity in two ways, namely (1) whether the dynamics of the model is stochastic, and (2) whether the cost takes stochasticity into consideration.

Previous studies have not dealt with either of these or considered only the first point. Previous studies that utilized control theory [5, 7, 9, 13, 12, 14, 15] applied the linear state-space model with deterministic dynamics and deterministic cost (Eq. (6)), while another study applied a stochastic model for brain dynamics [20] but calculated costs based on the deterministic framework.

To our knowledge, our present and recent studies [6] are the first to take account of stochasticity, not only with regard to brain dynamics but to cost. As we saw in Fig. 4, incorporating stochasticity in quantifying the control cost significantly changes the result. Thus, it will be desirable to assess the influence of stochasticity in the brain on control costs as we did in the present work.

### 4.2 Advantages of the Decomposition into Mean and Covariance Control Cost

One remarkable characteristic of the newly proposed cost is that it allows decomposition into mean and covariance control cost (Eq. (22)). A major theoretical advantage of this decomposition is that, as seen in Section 3.1.2, it enables us to separately examine the influence when taking covariance into account. Here, we discuss two significant aspects of this decomposition in neuroscience.

First, thanks to the decomposition, the stochastic cost can quantitatively compare the significance of contributions of two separate phenomena in a unified manner, namely the change in magnitude and co-variation of brain activities. Conventionally, changes in the magnitude of brain activities have been assessed through the estimated coefficients (often denoted as *β*) in the general linear model [36, 37]. The change in co-variation has been evaluated through the subtraction of correlation matrices [38, 39] or through network theoretic measures [40, 41, 42]. The dynamical change in co-variation (often referred to as functional connectivity in the field of neuroimaging) is called functional reconfiguration [43, 44, 45, 46]. In this way, classically, the change in magnitudes and the co-variation of activities have been examined separately in different contexts. In contrast, our use of decomposition has allowed us to quantitatively compare control costs for the mean and covariance from a unified perspective.

Second, the proposed framework enables us to investigate regions playing a significant role in the control of brain state transitions, as we have seen in Section 3.1.3. This topic is covered in the next section.

### 4.3 Significant Brain Regions in the Control of State Transitions

The present framework allows us to identify brain areas that contribute to controlling the mean and covariance separately. In Section 3.2.2, we saw the general tendencies of the mean and covariance input maps. Compared to the mean inputs, the covariance inputs are (1) more widespread (Fig. 5) and (2) less related to the regions of altered activities (Fig. 6). The first observation might reflect the fact that the alteration of covariance (termed functional reconfiguration [43, 44, 45, 46]) occurs all over the brain and thus broad input to control covariance may be necessary. The second observation means that we cannot estimate important regions for controlling covariance using only the magnitudes of activity. Examining the magnitudes of activity only may lead to the failure to identify significant regions controlling brain state transitions.

On examination of significant areas for control, an intriguing observation is that the posterior cingulate cortex (PCC) commonly contributes to controlling the covariance in many of the tasks examined (Fig. 7-a & b, Fig. 8, the middle panels). Among previous findings related to this result, the PCC is reported to be connected with many brain areas structurally [47] and functionally [48]. Although the PCC’s specific functions are not yet fully understood, previous studies have revealed that it is associated with many cognitive processes, including cognitive control [49], as shown using fMRI [50], and through recordings of single-cell firing rates [51]. From these studies, it is now considered that the PCC exerts influence on various regions and might thereby play a role in altering the functional connectivity of the brain [49, 50, 52]. Our finding that the PCC is significant in controlling covariance supports this view. Although this speculation needs further evidence, our study showed the importance of the PCC in changing functional connectivity from a control theoretic perspective.

Compared to the PCC and its contribution to controlling covariance, we found that the lower visual areas are included in the significant regions controlling both the mean and covariance (Fig. 7-a & b, Fig. 8, left panels). This might be due to the nature of the tasks recorded in the HCP dataset, where subjects were presented visual stimuli as part of the tasks.

### 4.4 Validity and Merits of the Linear Continuous Model for fMRI Data Analysis

In this study, we adopted the linear continuous-state model to describe brain dynamics in the fMRI data. It has been pointed out that a linear model may oversimplify the brain dynamics, which is complex and known to behave in nonlinear manners [53, 54, 55]. Although we acknowledge this limitation, a recent study has shown that the neural data, including the same fMRI data as we used in this study (HCP), can be better fitted with linear models than nonlinear ones, possibly because of temporal and spatial averaging effects [56]. Accordingly, this previous study [56] provides validity for our use of a linear model for the dataset.

To account for the nonlinearity of brain dynamics, another approach to modeling brain dynamics in the fMRI data is to use the discrete-space probabilistic model [6, 12, 57]. In our recent study [6], we used the same KL divergence as control cost in brain state transition as in the present study. The difference from our study lies in that in [6] we employed a different probabilistic model, where brain states are discrete.

Although the discrete model can incorporate nonlinearity, the linear model is more feasible than the discrete model in computing control costs in high-dimensional brain dynamics. In the discrete model, the computational cost of the control cost can easily explode, as the control cost is analytically intractable. We have to utilize iterative algorithms such as Sinkhorn’s algorithms to compute control costs [6, 58], and to lessen the computational burden of the algorithms, we have to coarse-grain and limit the number of brain states. In contrast, control cost in the linear continuous model is analytically tractable, enabling us to compute control costs in high-dimensional dynamics.

### 4.5 Future Directions

There are two possible extensions of our work: one is incorporating more general control inputs, and the other is the estimation of actual control costs from neural data.

For the first point, in this study, we considered limited situations of control where we implemented a model in which independent inputs are assigned to all nodes, as discussed in Section 2.1.1. In previous studies that have utilized the deterministic control theoretical framework in [5, 12, 13, 20], the model was grounded on more general cases wherein the system is described with an input term *Bu*, as in Eq. (2). We may be able to consider these general cases of the input matrix *B* in our stochastic framework with some additional techniques for computation (see [19] for details).

For the second point, we only considered the optimal control cost where brain state transitions are controlled in an optimal manner, with minimization of stochastic control cost. However, in real neural systems, state transitions are not controlled in an optimal manner. An intriguing future direction will be to compare the optimal and actual dynamics using neural data during tasks. Estimating the control cost from real time-series data requires the estimation of time-varying control inputs. This is generally a difficult estimation problem that requires more sophisticated techniques [59]. Quantifying the actual control cost in real neural data will provide another new insight into cognitive processes, which we plan to pursue in future research.

## Acknowledgement

Shunsuke Kamiya was supported by JSPS KAKENHI Grant Number JP22J23428. Masafumi Oizumi and Shuntaro Sasai were supported by JST Moonshot R&D Grant Number JPMJMS2012. Masafumi Oizumi was supported by JST CREST Grant Number JPMJCR1864, and JSPS KAKENHI Grant Numbers JP18H02713 and 20H05712.

---

|  | Previous framework | Proposed framework |
| --- | --- | --- |
|  | Controlled process<br>$\dot{x} = Ax + v$<br>Uncontrolled process<br>$\dot{x} = Ax$<br>$t = 0$ $t = T$ | Controlled process<br>$dx = Axd t + vdt + Cdw$<br>Uncontrolled process<br>$t = 0$ $t = T$ |
| Dynamics | Deterministic | **Stochastic process** |
| Transition | Point → point | **Prob. dist. → Prob. dist.** |
| Control cost | Squared integral<br>$\min_u \int_0^T \ u\ _2^2 dt.$ | **KL divergence**<br>$\int \log \frac{d\tilde{\mathcal{P}}}{d\mathcal{P}} d\tilde{\mathcal{P}}$ |

## References

[1] Breakspear, M. Dynamic models of large-scale brain activity. Nat. Neurosci. 20, 340–352 (2017).

[2] Kringelbach, M. L. & Deco, G. Brain states and transitions: Insights from computational neuroscience. Cell Rep. 32, 108128 (2020).

[3] McKenna, T. M., McMullen, T. A. & Shlesinger, M. F. The brain as a dynamic physical system. Neuroscience 60, 587–605 (1994).

[4] Botvinick, M. M. & Cohen, J. D. The computational and neural basis of cognitive control: charted territory and new frontiers. Cogn. Sci. 38, 1249–1285 (2014).

[5] Gu, S. et al. Controllability of structural brain networks. Nat. Commun. 6, 8414 (2015).

[6] Kawakita, G., Kamiya, S., Sasai, S., Kitazono, J. & Oizumi, M. Quantifying brain state transition cost via schr ö dinger bridge. Network Neuroscience 6, 118–134 (2022).

[7] Braun, U. et al. Brain network dynamics during working memory are modulated by dopamine and diminished in schizophrenia. Nat. Commun. 12, 3478 (2021).

[8] Deco, G. et al. Awakening: Predicting external stimulation to force transitions between different brain states. Proc. Natl. Acad. Sci. U. S. A. 116, 18088–18097 (2019).

[9] Szymula, K. P., Pasqualetti, F., Graybiel, A. M., Desrochers, T. M. & Bassett, D. S. Habit learning supported by efficiently controlled network dynamics in naive macaque monkeys (2020). 2006.14565.

[10] Liu, Y. Y., Slotine, J. J. & Barab ási, A. L. Controllability of complex networks. Nature 473, 167–173 (2011).

[11] Liu, Y. Y. & Barab ási, A. L. Control principles of complex systems. Rev. Mod. Phys. 88, 1–61 (2016).

[12] Cornblath, E. J. et al. Temporal sequences of brain activity at rest are constrained by white matter structure and modulated by cognitive demands. Commun Biol 3, 261 (2020).

[13] Kim, J. Z. et al. Role of graph architecture in controlling dynamical networks with applications to neural systems. Nat. Phys. 14, 91–98 (2018).

[14] Gu, S. et al. Optimal trajectories of brain state transitions. Neuroimage 148, 305–317 (2017).

[15] Stiso, J. et al. White matter network architecture guides direct electrical stimulation through optimal state transitions. Cell Rep. 28, 2554–2566.e7 (2019).

[16] Rieke, F. Spikes: exploring the neural code (MIT press, 1999).

[17] Faisal, A. A., Selen, L. P. J. & Wolpert, D. M. Noise in the nervous system. Nat. Rev. Neurosci. 9, 292–303 (2008).

[18] Van Essen, D. C. et al. The WU-Minn human connectome project: an overview. Neuroimage 80, 62–79 (2013).

[19] Kamiya, S., Kawakita, G., Sasai, S., Kitazono, J. & Oizumi, M. Supplementary information of optimal control costs of brain state transitions in linear stochastic systems. bioRxiv (2022).

[20] Deng, S. & Gu, S. Controllability analysis of functional brain networks (2020). 2003.08278.

[21] Rieke, F. Spikes: exploring the neural code (MIT press, 1999).

[22] Chen, Y., Georgiou, T. T. & Pavon, M. On the relation between optimal transport and schr ödinger bridges: A stochastic control viewpoint. J. Optim. Theory Appl. 169, 671–691 (2016).

[23] L éonard, C. A survey of the schrödinger problem and some of its connections with optimal transport (2013). 1308.0215.

[24] Dai Pra, P. A stochastic control approach to reciprocal diffusion processes. Appl. Math. Optim. 23, 313–329 (1991).

[25] Schrodinger, E. Uber die umkehrung der naturgesetze. sitz. ber. der preuss. Akad. Wissen., Berlin Phys. Math 144 (1931).

[26] Chen, Y., Georgiou, T. T. & Pavon, M. Optimal steering of a linear stochastic system to a final probability distribution, part I. IEEE Trans. Automat. Contr. 61, 1158–1169 (2016).

[27] Chen, Y., Georgiou, T. T. & Pavon, M. On the relation between optimal transport and schr ö dinger bridges: A stochastic control viewpoint. J. Optim. Theory Appl. 169, 671–691 (2016).

[28] Ito, T. et al. Task-evoked activity quenches neural correlations and variability across cortical areas. PLoS Comput. Biol. 16, e1007983 (2020).

[29] Satterthwaite, T. D. et al. An improved framework for confound regression and filtering for control of motion artifact in the preprocessing of resting-state functional connectivity data. Neuroimage 64, 240–256 (2013).

[30] Schaefer, A. et al. Local-Global parcellation of the human cerebral cortex from intrinsic functional connectivity MRI. Cereb. Cortex 28, 3095–3114 (2018).

[31] Poldrack, R. A., Mumford, J. A. & Nichols, T. E. Handbook of functional MRI data analysis (Cambridge University Press, 2011).

[32] Yan, G. et al. Spectrum of controlling and observing complex networks. Nature Physics 11, 779–786 (2015).

[33] Brunton, S. L. & Kutz, J. N. Data-driven science and engineering: Machine learning, dynamical systems, and control (Cambridge University Press, 2022).

[34] Parker Singleton, S. et al. LSD flattens the brain’s energy landscape: evidence from receptor-informed network control theory (2021).

[35] Van Essen, D. C. et al. The WU-Minn human connectome project: an overview. Neuroimage 80, 62–79 (2013).

[36] Friston, K. J. et al. Analysis of fmri time-series revisited. Neuroimage 2, 45–53 (1995).

[37] Ashby, F. G. Statistical analysis of fMRI data (MIT press, 2019).

[38] Cole, M. W., Bassett, D. S., Power, J. D., Braver, T. S. & Petersen, S. E. Intrinsic and task-evoked network architectures of the human brain. Neuron 83, 238–251 (2014).

[39] Cole, M. W., Ito, T., Bassett, D. S. & Schultz, D. H. Activity flow over resting-state networks shapes cognitive task activations. Nat. Neurosci. 19, 1718–1726 (2016).

[40] Finc, K. et al. Dynamic reconfiguration of functional brain networks during working memory training. Nature communications 11, 1–15 (2020).

[41] Hahn, A. et al. Reconfiguration of functional brain networks and metabolic cost converge during task performance. Elife 9, e52443 (2020).

[42] Braun, U. et al. Dynamic reconfiguration of frontal brain networks during executive cognition in humans. Proc. Natl. Acad. Sci. U. S. A. 112, 11678–11683 (2015).

[43] Kitzbichler, M. G., Henson, R. N. A., Smith, M. L., Nathan, P. J. & Bullmore, E. T. Cognitive effort drives workspace configuration of human brain functional networks. J. Neurosci. 31, 8259–8270 (2011).

[44] Davison, E. N. et al. Brain network adaptability across task states. PLoS Comput. Biol. 11, e1004029 (2015).

[45] Stitt, I. et al. Dynamic reconfiguration of cortical functional connectivity across brain states. Sci. Rep. 7, 8797 (2017).

[46] Krienen, F. M., Yeo, B. T. T. & Buckner, R. L. Reconfigurable task-dependent functional coupling modes cluster around a core functional architecture. Philos. Trans. R. Soc. Lond. B Biol. Sci. 369 (2014).

[47] Hagmann, P. et al. Mapping the structural core of human cerebral cortex. PLoS Biol. 6, e159 (2008).

[48] Leech, R., Braga, R. & Sharp, D. J. Echoes of the brain within the posterior cingulate cortex. Journal of Neuroscience 32, 215–222 (2012).

[49] Leech, R. & Sharp, D. J. The role of the posterior cingulate cortex in cognition and disease. Brain 137, 12–32 (2014).

[50] Lin, P. et al. Static and dynamic posterior cingulate cortex nodal topology of default mode network predicts attention task performance. Brain imaging and behavior 10, 212–225 (2016).

[51] Hayden, B., Smith, D. V. & Platt, M. Cognitive control signals in posterior cingulate cortex. Frontiers in human neuroscience 4, 223 (2010).

[52] Leech, R., Kamourieh, S., Beckmann, C. F. & Sharp, D. J. Fractionating the default mode network: distinct contributions of the ventral and dorsal posterior cingulate cortex to cognitive control. Journal of Neuroscience 31, 3217–3224 (2011).

[53] Rabinovich, M. I. & Muezzinoglu, M. Nonlinear dynamics of the brain: emotion and cognition. Physics-Uspekhi 53, 357 (2010).

[54] Freeman, W. J. Nonlinear gain mediating cortical stimulus-response relations. Biol. Cybern. 33, 237–247 (1979).

[55] Stephan, K. E. et al. Nonlinear dynamic causal models for fmri. Neuroimage 42, 649–662 (2008).

[56] Nozari, E. et al. Is the brain macroscopically linear? a system identification of resting state dynamics (2020).

[57] Lynn, C. W., Cornblath, E. J., Papadopoulos, L., Bertolero, M. A. & Bassett, D. S. Broken detailed balance and entropy production in the human brain. Proceedings of the National Academy of Sciences 118 (2021).

[58] Cuturi, M. Sinkhorn distances: Lightspeed computation of optimal transport. Advances in neural information processing systems 26 (2013).

[59] Milde, T. et al. Time-variant partial directed coherence in analysis of the cardiovascular system. a methodological study. Physiological measurement 32, 1787 (2011).

