## Supplementary Information for "Optimal Control Costs of Brain State Transitions in Linear Stochastic Systems"

### Supplementary Note 1 Control Energy in the Deterministic System

In the classical linear system described as (2), we consider minimizing the quadratic control cost

$$\min_u \int_0^T \|u\|_2^2 dt. \tag{SI-1}$$

The minimizing input $u^*(t)$ is known to exist with a controllable pair $(A, B)$, and the minimal value of the integral is expressed as

$$\mathcal{J}^*_{\mathrm{det}} := {}^{\mathrm{t}}(\mu_T - \Phi(T,0)\mu_0) M_{\mathrm{det}}(T,0)^{-1} (\mu_T - \Phi(T,0)\mu_0), \tag{SI-2}$$

where $\Phi$ is a state transition matrix such that

$$\partial_t \Phi(t,s) = A\Phi(t,s),\ \Phi(t,t) = I \ \Leftrightarrow\ \Phi(t,s) = e^{A(t-s)}, \tag{SI-3}$$

and $M$ is the control Gramian (which is nonsingular when the system is controllable),

$$M_{\mathrm{det}}(t,s) = \int_s^t \Phi(t,\tau) B\, {}^{\mathrm{t}}B\, {}^{\mathrm{t}}\Phi(t,\tau) d\tau. \tag{SI-4}$$

### Supplementary Note 2 Equivalence between KL cost and Quadratic Cost

In this section, we show that KL minimization boils down to an optimal control problem where the cost function is the expectation of the quadratic input with respect to controlled process [27, 24]. First, we observe the following two propositions.

**Proposition SI 2.1** (Chen [26]). The next two probability laws $\mathcal{Q}_1$ and $\mathcal{Q}_2$ are equal:

1. The probability law $\mathcal{Q}_1$ that minimizes the KL divergence:

   $$\mathcal{Q}_1 = \arg\min_{\tilde{\mathcal{P}}} D_{\mathrm{KL}}\left(\tilde{\mathcal{P}}, \mathcal{P}\right), \tag{SI-5}$$

   the boundary conditions given by Eq. (12).

2. The probability law $\mathcal{Q}_2$ induced by the next process:

$$dx(t) = Ax(t)dt + v^* \left(x(t), t\right) dt + Cdw(t) \quad \text{(SI-6)}$$

where $v^* : \mathbb{R}^n \times [0, T] \to \mathbb{R}^n$ is the process that minimizes the next functional of $v : \mathbb{R}^n \times [0, T] \to \mathbb{R}^n$,

$$\mathcal{J}(v) = \mathbb{E}_{\tilde{\mathcal{P}}} \left[ \int_0^T \|v\|^2_{(C\,{}^{\mathrm{t}}C)^{-1}} \, dt \right], \quad \text{(SI-7)}$$

where $\mathbb{E}_{\tilde{\mathcal{P}}}[\cdot]$ represents the expectation on a probability law $\tilde{\mathcal{P}}$ and $\|v\|^2_{(C\,{}^{\mathrm{t}}C)^{-1}} = {}^{\mathrm{t}}v \left(C\,{}^{\mathrm{t}}C\right)^{-1} v$. Here, $\tilde{\mathcal{P}}$ is a probability law induced by the next process:

$$dx(t) = Ax(t)dt + v \left(x(t), t\right) dt + Cdw(t). \quad \text{(SI-8)}$$

$v$ moves over the set such that the process (Eq. (SI-8)) satisfies the Gaussian boundary conditions given by Eq. (12) and

$$\mathbb{E}_{\tilde{\mathcal{P}}} \left[ \int_0^T \|v\|^2_{(C\,{}^{\mathrm{t}}C)^{-1}} \, dt \right] < \infty. \quad \text{(SI-9)}$$

*Proof.* See [26]. □

This proposition claims that the KL minimization problem is equivalent to the optimization problem where the cost function is a quadratic function (Eq. (SI-7)). The proposition also states that among numerous dynamics that satisfy the boundary conditions (Eq. (12)), we can confine our search space to the space that consists of the dynamics that can be described in the form of Eq. (SI-8) with an input term $v(x(t), t)$.

The above Proposition SI 2.1 tells nothing about how the values of the KL divergence (Eq. (SI-5)) and the quadratic cost (Eq. (SI-10)) are related. Surprisingly, the value of the KL divergence itself is equal to the quadratic cost function (Eq. (SI-10)), as shown in the next proposition.

**Proposition SI 2.2.** Let the uncontrolled process (Eq. (7)) and the controlled process (Eq. (SI-8)) admit solutions, and $\mathcal{P}$ and $\tilde{\mathcal{P}}$ probability distributions induced by the uncontrolled and controlled process, respectively. We assume that the two processes have the same initial distribution

$\pi_0$ (i.e., the probability distribution of $x(0)$) that is absolutely continuous with respect to the Lebesgue measure. If Eq. (SI-9) stands, then

$$D_{\mathrm{KL}}\left(\tilde{\mathcal{P}},\mathcal{P}\right)=\frac{1}{2}\mathbb{E}_{\tilde{\mathcal{P}}}\left[\int_0^T \|v\|^2_{(C^{t}C)^{-1}}\,dt\right]. \tag{SI-10}$$

*Proof.* We make use of the Girsanov theorem, which is a rule for transforming measures of processes expressed with different Brownian motions. This proof is based on [60] and [61]. Since $C$ is nonsingular, the controlled process (Eq. (SI-8)) is described as

$$d\tilde{x}(t)=C^{-1}AC\tilde{x}(t)dt+C^{-1}vdt+d\tilde{w}(t), \tag{SI-11}$$

where $\tilde{x}=C^{-1}x$. We denote the $\tilde{\mathcal{P}}$-Brownian motion by $\tilde{w}$ for consistency of notation. By Maruyama-Girsanov's theorem, the process $\tilde{x}$ can be described as a form of

$$d\tilde{x}(t)=C^{-1}AC\tilde{x}(t)dt+dw'(t), \tag{SI-12}$$

where $w'(t)$ is a $\mathcal{P}'$-Brownian motion. The probability law $\mathcal{P}'$ is given as , by Maruyama-Girsanov's theorem,

$$\frac{d\mathcal{P}'}{d\tilde{\mathcal{P}}}=\frac{d\pi_0(x)/dx}{d\pi_0(x)/dx}X(T)=X(T). \tag{SI-13}$$

Here, $X(t)$ is

$$X(t)=\exp\left(-\int_0^t\left(C^{-1}v(x(s),s)\right)dw'(s)-\frac{1}{2}\int_0^t\|C^{-1}v(x(s),s)\|^2ds\right), \tag{SI-14}$$

and is a martingale since $C^{-1}v$ satisfies Novikov condition by Eq. (SI-9). Thus,

$$\begin{aligned}D_{\mathrm{KL}}\left(\tilde{\mathcal{P}},\mathcal{P}\right)&=\int_S\log\frac{d\tilde{\mathcal{P}}}{d\mathcal{P}'}d\tilde{\mathcal{P}}\\
&=\frac{1}{2}\mathbb{E}_{\tilde{\mathcal{P}}}\left[\int_0^T C^{-1}v(x(s),s)dw'(s)+\int_0^T\|C^{-1}v(x(s),s)\|^2ds\right]\\
&=\frac{1}{2}\mathbb{E}_{\tilde{\mathcal{P}}}\left[\int_0^T\|C^{-1}v(x(s),s)\|^2ds\right].\end{aligned} \tag{SI-15}$$

The last equality arises from the fact that the expectation of the Ito integral is always zero. □

### Supplementary Note 3 Derivation of the stochastic control cost

In this section, we derive the analytical expression of Eq. (21) by answering the next problem. The problem statement and the derivation of $\mathcal{J}$ until Eq. (SI-30) are based on [26].

**Problem.** Consider the control statistics $v : \mathbb{R}^n \times [0, T] \to \mathbb{R}^n$ in the next $n$-dimensional dynamics

$$dx(t) = Ax(t)dt + v(t)dt + Cdw(t) \tag{SI-16}$$

where $w(t)$ is a standard $n$-dimensional normal Brownian motion, and $A \in \mathbb{R}^{n\times n},\ C \in \mathbb{R}^{n\times n}$. We assume $C$ to be nonsingular. The marginal distributions are bounded with

$$x(0) \sim \mathcal{N}(\mu_0, \Sigma_0), \quad x(T) \sim \mathcal{N}(\mu_T, \Sigma_T). \tag{SI-17}$$

Out of many possible $v(t)$, find one that minimizes the next cost function:

$$\mathcal{J}(v) = \mathbb{E}\left[\int_0^T \|v\|^2_{(C\,{}^{t}C)^{-1}} dt\right], \tag{SI-18}$$

when this value is finite.

**Solution.** In [26], the existence of the optimizer $v^*$ of Eq. (SI-7) under Eq. (SI-8) and Eq. (SI-9) was proved. The optimizer has a closed form solution,

$$v^*(x(t), t) = -C\,{}^{t}C\Pi(t)x(t) + C\,{}^{t}C\,{}^{t}m(t). \tag{SI-19}$$

Here, $\Pi(t)$ is the nonsingular solution of the next matrix Riccati equation

$$\dot{\Pi}(t) = -{}^{t}A\Pi(t) - \Pi(t)A + \Pi(t)C\,{}^{t}C\Pi(t), \tag{SI-20}$$

under

$$\begin{aligned}\Pi(0) = \Sigma_0^{-\frac{1}{2}} \Bigg(&\frac{1}{2}I + \Sigma_0^{\frac{1}{2}}\,{}^{t}\Phi(T,0)M(T,0)^{-1}\Phi(T,0)\Sigma_0^{\frac{1}{2}} \\ &- \left(\frac{1}{4}I + \Sigma_0^{\frac{1}{2}}\,{}^{t}\Phi(T,0)M(T,0)^{-1}\Sigma_T M(T,0)^{-1}\Phi(T,0)\Sigma_0^{\frac{1}{2}}\right)^{\frac{1}{2}}\Bigg)\Sigma_0^{-\frac{1}{2}},\end{aligned} \tag{SI-21}$$

and

$$\partial_t \Phi(t,s) = A\Phi(t,s),\ \Phi(t,t) = I \ \Leftrightarrow\ \Phi(t,s) = e^{A(t-s)}, \tag{SI-22}$$

$$\partial_t \Psi(t,s) = \left(A - C\,{}^t C \Pi(t)\right) \Psi(t,s), \quad \Psi(t,t) = I, \tag{SI-23}$$

$$M(t,s) = \int_s^t \Phi(t,\tau) C\,{}^t C\,{}^t\Phi(t,\tau) d\tau, \tag{SI-24}$$

$$G(t,s) = \int_s^t \Psi(t,\tau) C\,{}^t C^t \Psi(t,\tau) d\tau, \tag{SI-25}$$

$$m(t) = {}^t\Psi(T,t) G(T,0)^{-1} (\mu_T - \Psi(T,0)\mu_0). \tag{SI-26}$$

See [26] for the derivations. $G(T,0)$ is denoted by $G(T)$ for abbreviation.

Consider adding some terms to $\mathcal{J}$ to have

$$\begin{aligned}
\tilde{\mathcal{J}} &= \mathbb{E}\left[\int_0^T \|u(t)\|^2\ dt + {}^t x(T)\Pi(T)x(T) - 2\,{}^t m(T) x_T - {}^t x(0)\Pi(0)x(0) + 2\,{}^t m(0) x_0\right] \\
&= \mathbb{E}\left[\int_0^T \left(\|u(t)\|^2 + \left[{}^t x(t)\Pi(t)x(t) - 2\,{}^t m(t)x(t)\right]_0^T\right) dt\right],
\end{aligned} \tag{SI-27}$$

where $u(t) = C^{-1}v(t)$. Let a function $V(t)$ as

$$V(t) = {}^t x(t)\Pi(t)x(t) - 2\,{}^t m(t)x(t). \tag{SI-28}$$

By Ito's formula,

$$\begin{aligned}
dV &= \left({}^t x(\dot{\Pi} - 2\,{}^t\dot{m})x + \left({}^t x\,{}^t A + {}^t u\,{}^t C\right)(2\Pi x - 2m) + \frac{1}{2}\operatorname{tr}\left(2\,{}^t C\Pi C\right)\right) dt + {}^t C(2\Pi x - 2m) dw \\
&= \left({}^t x(\dot{\Pi} + {}^t A\Pi + \Pi A)x - 2({}^t\dot{m} + {}^t m A - {}^t u\,{}^t C\Pi)x - {}^t u\,{}^t C m + \operatorname{tr}({}^t C\Pi C)\right) dt + 2({}^t x\Pi - {}^t m)C dw \\
&= \left({}^t x\Pi C\,{}^t C\Pi x - 2({}^t m C\,{}^t C - {}^t u\,{}^t C)\Pi x - 2\,{}^t u\,{}^t C m + \operatorname{tr}({}^t C\Pi C)\right) dt + 2({}^t x\Pi - {}^t m)C dw.
\end{aligned} \tag{SI-29}$$

Substitute in Eq. (SI-27) to get

$$\begin{aligned}
\tilde{\mathcal{J}} &= \mathbb{E}\left[\int_0^T \left(\|u(t)\|^2 + {}^t x\Pi C\,{}^t C\Pi x - 2({}^t m C\,{}^t C - {}^t u\,{}^t C)\Pi x - 2\,{}^t u\,{}^t C m + \operatorname{tr}({}^t C\Pi C)\right) dt \right. \\
&\qquad\qquad \left. + 2\int_0^T ({}^t x\Pi - {}^t m)C\ dw\right] \\
&= \mathbb{E}\left[\|u + {}^t C\Pi x - {}^t C m\|^2 + \int_0^T \left(\operatorname{tr}({}^t C\Pi C) - {}^t m C\,{}^t C m\right) dt\right].
\end{aligned} \tag{SI-30}$$

Since $u^*(t) = C^{-1}v^*(t) = -{}^{\mathrm{t}}C\Pi x + {}^{\mathrm{t}}Cm(t)$, under $u(t) = u^*(t)$ we obtain

$$\tilde{\mathcal{J}}^* := \mathbb{E}\left[\int_0^T \left(\mathrm{tr}({}^{\mathrm{t}}C\Pi C) - {}^{\mathrm{t}}mC\,{}^{\mathrm{t}}Cm\right) dt\right]. \tag{SI-31}$$

The $\cdot^*$ denotes the value when $u(t)$ ($v(t)$) is optimal.

Also, with $\mu_d = \mu_T - \Psi(T,0)\mu_0$,

$$\begin{aligned}\int_0^T {}^{\mathrm{t}}mC\,{}^{\mathrm{t}}Cmdt &= \int_0^T \|{}^{\mathrm{t}}Cm\|^2 dt = \int_0^T \|{}^{\mathrm{t}}C\,{}^{\mathrm{t}}\Psi(T,t)G^{-1}\mu_d\|^2 dt \\ &= {}^{\mathrm{t}}\mu_d\,{}^{\mathrm{t}}G(T)^{-1}\underbrace{\int_0^T \Psi(T,t)C\,{}^{\mathrm{t}}C\,{}^{\mathrm{t}}\Psi(T,t)dt}_{G(T)}\,G(T)^{-1}\mu_d \\ &= {}^{\mathrm{t}}\mu_d G(T)^{-1}\mu_d,\end{aligned} \tag{SI-32}$$

since $G(T) = {}^{\mathrm{t}}G(T)$. By the definition of $\tilde{\mathcal{J}}$, when $u(t)$ ($v(t)$) is optimal,

$$\begin{aligned}\tilde{\mathcal{J}}^* - \mathcal{J}^* &= \mathbb{E}\left[{}^{\mathrm{t}}x(T)\Pi(T)x(T) - 2\,{}^{\mathrm{t}}\mu_T x(T) - {}^{\mathrm{t}}x(0)\Pi(0)x(0) + 2\,{}^{\mathrm{t}}\mu_0 x(0)\right] \\ &= \mathbb{E}\left[\mathrm{tr}(\Pi(T)x(T)\,{}^{\mathrm{t}}x(T)) - \mathrm{tr}(\Pi(0)x(0)\,{}^{\mathrm{t}}x(0))\right] - 2\,{}^{\mathrm{t}}m(T)\mathbb{E}[x(T)] + 2\,{}^{\mathrm{t}}m(0)\mathbb{E}[x(0)] \\ &= \mathrm{tr}(\Pi(T)(\Sigma_T + {}^{\mathrm{t}}\mu_T\mu_T) - 2\,{}^{\mathrm{t}}m(T)\mu_T - \mathrm{tr}(\Pi(0)(\Sigma_0 + {}^{\mathrm{t}}\mu_0\mu_0)) + 2\,{}^{\mathrm{t}}\mu_0\mu_0 \\ &= \mathrm{tr}\left(\Pi(T)(\Sigma_T + {}^{\mathrm{t}}\mu_T\mu_T) - \Pi(0)(\Sigma_0 + {}^{\mathrm{t}}\mu_0\mu_0)\right) - 2\,{}^{\mathrm{t}}\mu_d G(T)^{-1}\mu_d.\end{aligned} \tag{SI-33}$$

By Eq. (SI-27) and Eq. (SI-30),

$$\begin{aligned}\mathcal{J}^* = &\int_0^T \mathrm{tr}(\Pi C\,{}^{\mathrm{t}}C)dt - {}^{\mathrm{t}}\mu_d G(T)^{-1}\mu_d \\ &- \left(\mathrm{tr}\left(\Pi(T)(\Sigma_T + {}^{\mathrm{t}}\mu_T\mu_T) - \Pi(0)(\Sigma_0 + {}^{\mathrm{t}}\mu_0\mu_0)\right) - 2\,{}^{\mathrm{t}}\mu_d G(T)^{-1}\mu_d\right) \\ = &\int_0^T \mathrm{tr}(\Pi C\,{}^{\mathrm{t}}C)dt + {}^{\mathrm{t}}\mu_d G(T)^{-1}\mu_d \\ &- \mathrm{tr}\left(\Pi(T)(\Sigma_T + {}^{\mathrm{t}}\mu_T\mu_T) - \Pi(0)(\Sigma_0 + {}^{\mathrm{t}}\mu_0\mu_0)\right).\end{aligned} \tag{SI-34}$$

### Supplementary Note 4 Correspondence of the Mean Control Cost and the Deterministic Control Cost

As explained in the main text, there is a clear correspondence between the mean control cost $\mathcal{J}_\mu^*$ and the deterministic control cost $\mathcal{J}_{\mathrm{det}}$. To see this correspondence, we need a special condition that the input matrix $B$ in the deterministic equation (Eq. (2)) be equal to the diffusion matrix

$C$.

**Theorem SI 4.1.** *The control cost necessary for steering the mean, and the control cost defined in the deterministic systems, $\mathcal{J}_{det}$, are identical, namely,*

$$\mathcal{J}_\mu^* = \mathcal{J}_{det}^*, \tag{SI-35}$$

*where $\mathcal{J}_\mu^*$ is given as Eq. (23), and $\mathcal{J}_{det}^*$ is the (deterministic) control cost of the next linear invariant system,*

$$\dot{x}(t) = Ax(t) + Cu(t), \qquad x(0) = \mu_0,\ x(T) = \mu_T, \tag{SI-36}$$

*where $A, C \in \mathbb{R}^{n\times n}$, $\mathrm{rank}(C) = n$, and by Eq. (SI-2), written as*

$$\mathcal{J}_{det}^* = {}^{t}(\mu_T - \Phi(T,0)\mu_0)M(T)^{-1}(\mu_T - \Phi(T,0)\mu_0). \tag{SI-37}$$

Before proceeding to the proof, we start by showing the next lemma.

**Lemma SI 4.2.** *The next equality stands:*

$$\forall t \in [0,T] \quad G(t) - Q(t) = -\Psi(t,0)Q_0\,{}^{t}\Psi(t,0). \tag{SI-38}$$

*$\Psi$ and $G$ are defined as in Eq. (SI-23) and Eq. (SI-25), and $Q(t) = \Pi(t)^{-1}$ where $\Pi$ is defined as in Eq. (SI-20) and Eq. (SI-21).*

*Proof.* $\Pi(t)$ can also made to be always nonsingular in $\forall t \in [0,T]$ [26]. Let $\tilde{G} : [0,T] \to \mathbb{R}^{n\times n}$ be

$$\tilde{G}(t) = Q(t) - \Psi(t,0)Q_0\,{}^{t}\Psi(t,0). \tag{SI-39}$$

One has

$$\begin{aligned}\partial_t\tilde{G} &= \partial_t Q - (A - S\Pi(t))\Psi(t,0)Q_0\,{}^{t}\Psi(t,0) - \Psi(t,0)Q_0\,{}^{t}\Psi(t,0)\,{}^{t}(A - S\Pi(t))\\ &= AQ + Q\,{}^{t}A - S + (A - S\Pi(t))(\tilde{G} - Q) + (\tilde{G} - Q)\,{}^{t}(A - S\Pi(t))\\ &= (A - S\Pi(t))(\tilde{G} - Q) + (\tilde{G} - Q)\,{}^{t}(A - S\Pi(t)) + S.\end{aligned} \tag{SI-40}$$

This is equivalent to the dynamics that $G(t)$ follows, as by Eq. (SI-25) and the Leibniz rule one obtains

$$\partial_t G(t) = \partial_t \int_0^t \Psi(t,\tau) S\, {}^{t}\Psi(t,\tau) d\tau$$
$$= \Psi(t,t) S\, {}^{t}\Psi(t,t) + \int_0^t \partial_t \left(\Psi(t,\tau) S\, {}^{t}\Psi(t,\tau)\right) d\tau$$
$$= (A - S\Pi(t))(G - Q) + (G - Q)\, {}^{t}(A - S\Pi(t)) + S.$$

And $G(0) = \tilde{G}(0) = 0$ by simple substitution. Former studies including [62] have guaranteed the existence and uniqueness of the solution of the linear Lyapunov equation, thus

$$\tilde{G}(t) = G(t). \tag{SI-41}$$

□

*Proof of the theorem.* By plugging in the expression of $\mu_d$, one obtains

$$\mathcal{J}_\mu^* = {}^{t}\mu_T (G(T)^{-1} - \Pi(T))\mu_T - 2\, {}^{t}\mu_T G(T)^{-1} \Psi(T,0)\mu_0 + {}^{t}\mu_0 ({}^{t}\Psi(T,0) G(T)^{-1} \Psi(T,0) + \Pi(0))\mu_0$$
$$= {}^{t}\mu_T (G(T)^{-1} - \Pi(T))\mu_T - 2\, {}^{t}\mu_T G(T)^{-1} \Psi(T,0)\mu_0 + {}^{t}\mu_0 ({}^{t}L(0)^{-1} + \Pi(0))\mu_0, \tag{SI-42}$$
$$\mathcal{J}_{\text{det}}^* = {}^{t}\mu_T M(T)^{-1} \mu_T - 2\, {}^{t}\mu_T M(t)^{-1} \Phi(T,0)\mu_0 + {}^{t}\mu_0 L(0)^{-1}\mu_0, \tag{SI-43}$$

where one denotes, for abbreviation,

$$M(t) = M(t,0),\ G(t) = G(t,0), \tag{SI-44}$$
$$N(t) = N(T,t),\ N(t,s) = \int_s^t \Phi(s,\tau) C\, {}^{t}C\, {}^{t}\Phi(s,\tau)\ d\tau, \tag{SI-45}$$
$$L(t) = L(T,t),\ L(t,s) = \int_s^t \Psi(t,\tau) C\, {}^{t}C\, {}^{t}\Psi(t,\tau)\ d\tau. \tag{SI-46}$$

Note that $M(t), N(t)$, and $L(t)$ are all supposed to be nonsingular by the controllability condition. $\Pi(t)$ can also made to be always nonsingular in $\forall t \in [0,T]$ [26] and we let $Q(t) = \Pi(t)^{-1}$. To prove $\mathcal{J}_\mu^* = \mathcal{J}_{\text{det}}^*$, it is sufficient to show

$$M(T)^{-1} = G(T)^{-1} - \Pi(T), \tag{SI-47}$$
$$N(0)^{-1} = L(0)^{-1} + \Pi(0), \tag{SI-48}$$
$$M(T)^{-1}\Phi(T,0) = G(T)^{-1}\Psi(T,0). \tag{SI-49}$$

In fact, the equalities Eq. (SI-47) to Eq. (SI-49) do not only stand at $t = T$ but in $\forall t \in (0, T]$, namely

$$M(t)^{-1} = G(t)^{-1} - \Pi(t) \quad \forall t \in (0, T], \tag{SI-50}$$

$$N(t)^{-1} = L(t)^{-1} + \Pi(t) \forall t \in [0, T), \tag{SI-51}$$

$$M(t)^{-1}\Phi(t, 0) = G(t)^{-1}\Psi(t, 0), \quad \forall t \in (0, T]. \tag{SI-52}$$

We begin by showing Eq. (SI-50). By Lemma SI 4.2, $G - Q$ is always invertible in $t \in [0, T]$, as $\Psi(t, 0)$ and $Q(0)$ are invertible in the same range of $t$. Let $\xi : [0, T] \to \mathbb{R}^{n\times n}$ as

$$\xi(t) = -Q(t) \left(G(t) - Q(t)\right)^{-1} G(t) = \left(G(t)^{-1} - \Pi(t)\right)^{-1}. \tag{SI-53}$$

Differentiating this with $t$ and substitutions gives

$$\partial_t \xi = A\xi + \xi^{\,\mathrm{t}}A + S. \tag{SI-54}$$

Getting back to Eq. (SI-47), $M(t)$ satisfies

$$\partial_t M = AM + M^{\,\mathrm{t}}A + S, \tag{SI-55}$$

which is identical to the dynamics of $\xi$. The existence and uniqueness are guarantied in Lyapunov equations and by $\xi(0) = M(0) = 0$,

$$\xi(t) = M(t) \tag{SI-56}$$

in all $t \in [0, T]$. Thus $\xi^{-1}(t) = M^{-1}(t)$ for $t \in (0, T]$ stands.

One can prove the equality Eq. (SI-51) quite in the same manner.

As for Eq. (SI-52), let $\eta : [0, T] \to \mathbb{R}^{n\times n}$ as

$$\eta = (E - G(t)\Pi(t))\Phi(t, 0) \tag{SI-57}$$

where $E$ is the $n$-dimensional identity matrix. Now

$$\begin{aligned} \partial_t \eta = -\partial_t(G\Pi)\Phi(t, 0) + \eta\partial_t\Phi(t, 0) = (A - S\Pi)(I - G\Pi)\Psi(t, 0) \\ = (A - S\Pi)\eta. \end{aligned} \tag{SI-58}$$

Then $\eta$ satisfies the same dynamics as $\Psi(t,0)$, where by definition

$$\partial_t \Psi(t,0) = (A - S\Pi)\Psi(t,0). \tag{SI-59}$$

Also $\eta(0) = \Psi(0,0) = E$ holds, which concludes $\eta(t) = \Psi(t,0)$ in all $t \in [0,T]$. Then when $t > 0$,

$$G(t)(G(t)^{-1} - \Pi)\Phi(t,0) = \Psi(t,0) \tag{SI-60}$$

holds, which implies

$$(G^{-1} - \Pi)\Phi(t,0) = G(t)^{-1}\Psi(t,0). \tag{SI-61}$$

By Eq. (SI-47),

$$M(t)^{-1}\Phi(t,0) = G(t)^{-1}\Psi(t,0). \tag{SI-62}$$

□

### Supplementary Note 5 Derivation of (29)

Using Fubini's theorem,

$$\begin{aligned}
&\mathbb{E}\left[\int_0^T \|v_k^*\|^2 dt\right] = \mathbb{E}\left[\int_0^T (v^* {}^{t}v^*)_{kk} dt\right] \\
&= \left(\mathbb{E}\left[\int_0^T v^* {}^{t}v^* dt\right]\right)_{kk} \\
&= \left(C\ \mathbb{E}\left[\int_0^T C^{-1}v^* {}^{t}(C^{-1}v^*)dt\right] {}^{t}C\right)_{kk} \\
&= \left(C {}^{t}C\mathbb{E}\left[\int_0^T (\Pi(t)x(t) - m(t))\, {}^{t}(\Pi(t)x(t) - m(t))\, dt\right] C {}^{t}C\right)_{kk} \qquad \text{(by Eq. (SI-19))} \\
&= \left(S_C\mathbb{E}\left[\int_0^T \left(\Pi x {}^{t}x\Pi - 2m {}^{t}x\Pi + m {}^{t}m\right) dt\right] S_C\right)_{kk} \\
&= \left(S_C \int_0^T \mathbb{E}\left[\left(\Pi x {}^{t}x\Pi - 2m {}^{t}x\Pi + m {}^{t}m\right)\right] dt S_C\right)_{kk} \\
&= \left(S_C \int_0^T \left(\Pi\left(\Sigma(t) + \mu(t) {}^{t}\mu(t)\right)\Pi - 2m {}^{t}\mu(t)\Pi + m {}^{t}m\right) dt\ S_C\right)_{kk}.
\end{aligned}$$

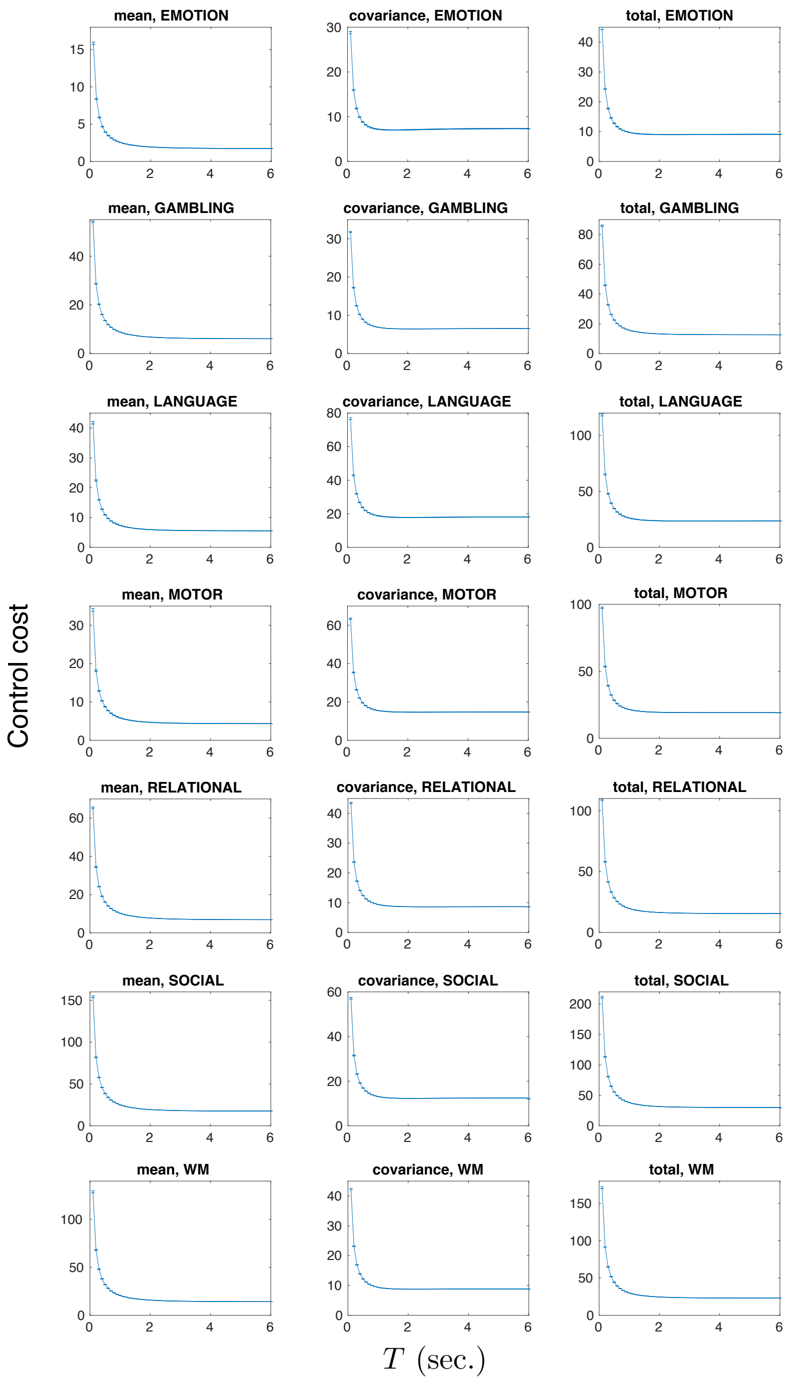


**Supplementary Figure 1: Stochastic control cost values of all seven tasks with $T$ ranging from** $0.1$ **to** $5.0$ **seconds.** The $T$ values were computed in 0.1-second increments. For each task: left panel, the mean control costs; middle, the covariance control costs; and right, the total control costs. The time range was based on our assumption that control of state transition occurs in a relatively short time, not more than an order of $\times 10^0$ seconds. We observed that the control cost values change according to $T$ for all tasks. Specifically, all three types of cost (the mean, covariance, and total control costs) rapidly decrease when $T$ increases and converge to a fixed value when $T$ is large enough. All the curves seem to plateau at approximately $T = 1.0$ seconds. We therefore chose $T = 1.0$ as representative in the main text. The ratios and the ordering of the seven tasks, which are examined in Section 3.2.1, did not change drastically with various $T$ values.

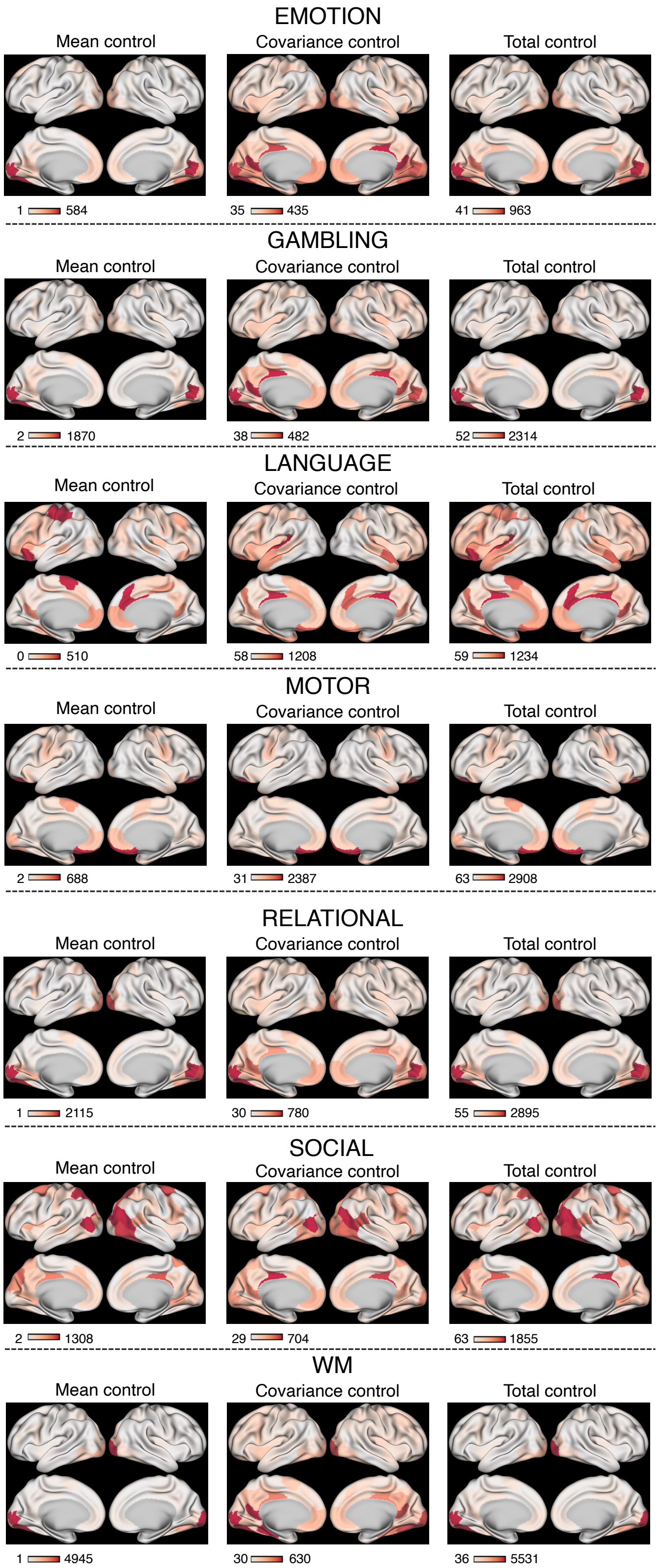


**Supplementary Figure 2: Inputs at each brain region of the seven tasks in the HCP dataset.** For each task: left panel, the mean input map; middle, the covariance input map; and right, the total input map. Color bars represent the quantity of control inputs ($\times 10^{-5}$).

### Supplementary Note 6 Limitations of the theoretical framework

In this study, we considered limited situations of control wherein we implemented a model in which independent inputs are assigned to all nodes. This is mathematically denoted as a stochastic system with an input term $v : [0, T] \to \mathbb{R}^n$ (equation (SI-8)). In the standard control theoretic framework, the input term $v$ is often described as $Bu$, where $B \in \mathbb{R}^{n\times m}$ is an input matrix and $u : [0, T] \to \mathbb{R}^m$ is the control input (see equation (2)). We did not incorporate the input matrix $B$, which means that we only consider where $B \in \mathbb{R}^{n\times n}$ is an identity matrix.

In previous studies that have utilized the deterministic control theoretical framework, the model was grounded on more general cases wherein the system is described with an input term $Bu$, as in equation (2). For example, in [12, 13, 20], $B \in \mathbb{R}^{n\times m}$, $m \neq n$ was set so that it represents independent inputs in a subgroup of nodes $\{k_1, \cdots, k_m\}$,

$$(B)_{ij} = \delta_{k_j j} \quad (i = 1, \cdots, n,\ j = 1, \cdots, m), \tag{SI-63}$$

where $\delta$ is the Kronecker delta. These studies parametrized $m$, the number of input nodes, to consider the best control strategy. Another study [5] modeled the input nodes to only one, and set $B$ as an $n$-by-1 matrix (or an $n$-dimensional vector),

$$(B)_i = \begin{cases} 1 & (i = k) \\ 0 & (\text{otherwise}) \end{cases}, \tag{SI-64}$$

for a fixed $k \in \{1, \cdots, n\}$.

Even though we can technically consider such general cases where the input matrix $B$ satisfies $\in \mathbb{R}^{n\times m}$ $(n \neq m)$ in our stochastic framework, computing the control costs in such models is more difficult both analytically and numerically. According to a previous study [63], if $B$ is rank deficient, we cannot find an analytical solution and hence have to resort to a numerical convex optimization with time discretization, which takes enormous computational time.

To alleviate computational time in high-dimensional data, we may need different approaches, such as dimensionality reduction. It has been reported that the major neural activity patterns are confined on a low dimensional space, named neural manifolds [64, 65, 66, 67]. Based on these findings, we may first apply the dimensionality reduction to high-dimensional neural data and then fit the model with a general input matrix $B$.
